# Steatotic liver disease induced by TCPOBOP-activated hepatic constitutive androstane receptor: Primary and secondary gene responses with links to disease progression

**DOI:** 10.1101/2024.02.16.580717

**Authors:** Ravi Sonkar, Hong Ma, David J. Waxman

## Abstract

Constitutive Androstane Receptor (CAR, *Nr1i3*), a liver nuclear receptor and xenobiotic sensor, induces drug, steroid and lipid metabolizing enzymes, stimulates liver hypertrophy and hyperplasia, and ultimately, hepatocellular carcinogenesis. The mechanisms linking early CAR responses to subsequent disease development are poorly understood. Here we show that exposure of CD-1 mice to TCPOBOP, a halogenated xenochemical and selective CAR agonist ligand, induces pericentral steatosis marked by hepatic accumulation of cholesterol and neutral lipid, and elevated circulating alanine aminotransferase levels, indicating hepatocyte damage. TCPOBOP-induced steatosis was weaker in the pericentral region but stronger in the periportal region in females compared to males. Early (1-day) TCPOBOP transcriptional responses were enriched for CAR-bound primary response genes, and for lipid and xenobiotic metabolism and oxidative stress protection pathways; late (2-wk) TCPOBOP responses included many CAR binding-independent secondary response genes, with enrichment for immune response, macrophage activation, and cytokine and reactive oxygen species production. Late upstream regulators specific to TCPOBOP-exposed male liver were linked to pro-inflammatory responses and hepatocellular carcinoma progression. TCPOBOP administered weekly to male mice using a high corn oil vehicle activated carbohydrate-responsive transcription factor (MLXIPL)-regulated target genes, dysregulated mitochondrial respiratory and translation regulatory pathways, and induced more advanced liver pathology. Thus, TCPOBOP exposure recapitulates histological and gene expression changes characteristic of emerging steatotic liver disease, including secondary expression changes in liver non-parenchymal cells indicative of transition to a more advanced disease state. Upstream regulators of both the early and late TCPOBOP gene responses include novel biomarkers for foreign chemical-induced metabolic dysfunction-associated steatotic liver disease.

## Introduction

Metabolic dysfunction-associated steatotic liver disease (MASLD), previously referred to as non-alcoholic fatty liver disease, is a widespread chronic liver disease associated with a lipotoxic environment that results from the pathological accumulation of triglycerides in hepatocytes, termed hepatic steatosis [1]. The early stages of MASLD, involving simple steatosis, can progress to metabolic dysfunction-associated steatohepatitis (MASH), which is characterized by hepatic inflammation and fibrosis and may progress to liver cirrhosis and necrosis [2, 3]. MASLD increases the risk for developing coronary heart disease and type 2 diabetes and is a leading cause of hepatocellular carcinoma (HCC) [4]. Sex differences characterize steatotic liver disease, with the prevalence and severity of MASLD, and its progression to MASH, liver cirrhosis and HCC being greater in males and in post-menopausal females than premenopausal females [5–7]. The molecular mechanisms underlying the development of MASLD and its progression to MASH and HCC are only partially understood, with important recent advances coming from genetic studies and global transcriptomic analyses in high fat dietary exposure models of disease [8–11].

Environmental chemicals and toxicants, including many persistent organic pollutants and other endocrine-disrupting chemicals, have long been associated with the development of steatotic liver disease [12–14]. The underlying mechanisms of toxicant-associated liver disease [15, 16] are likely to be multifactorial, given the wide spectrum of chemical exposures that can induce these liver pathologies. Many environmental chemicals that induce MASLD are ligands for transcription factors from the Nuclear Receptor (NR) gene superfamily, most notably CAR (constitutive androstane receptor, Nr1i3), PXR (pregnane X receptor, Nr1i2) and PPARA, which regulate genes of xenobiotic and energy metabolism by pathways that can either contribute to or protect from MASLD development [17]. Many foreign chemicals have broad specificities for receptor activation, which enables them to activate multiple receptors, including Ahr (aryl hydrocarbon receptor) [18], by either direct or indirect mechanisms, which further complicates efforts to elucidate underlying mechanisms of action.

CAR can be activated by structurally diverse drugs and environmental chemicals [19, 20], resulting in increased transcription of genes for many drug-metabolizing enzymes, including phase-I P450 enzymes, phase-II UDP-glucuronosyltransferases and sulfotransferases, as well as transporters active in drug uptake and efflux [21]. CAR also has important effects on endogenous lipid and energy metabolism [22, 23] and can lessen the hepato-steatotic effects of high fat dietary exposures [24, 25]. In the inactive, unliganded state, CAR is phosphorylated at threonine-38 and retained in the cytoplasm in a protein complex with heat shock protein 90 and cytoplasmic CAR retention protein [26]. CAR can be activated either by binding an agonist ligand [27], or indirectly, via a signaling pathway induced by non-ligand CAR agonists, such as phenobarbital [28]. Both activation mechanisms dephosphorylate CAR at threonine-38 and stimulate translocation of CAR to the nucleus [26], where it heterodimerizes with RXR and binds to enhancer sequences linked to the transcriptional activation of CAR target genes [29–31].

TCPOBOP (1,4-bis[2-(3,5-dichloropyridyloxy)]benzene) is an agonist ligand [27] that is highly selective for CAR [32]. TCPOBOP can therefore be used to study the effects of xenobiotic-activated CAR without the complexities that arise with polychlorinated biphenyls and other foreign chemicals, many of which activate and/or alter the expression levels of Ahr and/or other NR superfamily members [33–36]. TCPOBOP induces CAR-dependent hepatomegaly, leading to a substantial increase in liver size within a few days due to short-term induction of hepatocyte proliferation combined with hepatocellular hypertrophy [37–39]. TCPOBOP exposure induces widespread effects on the liver transcriptome, which have been characterized as early as 3 h [40, 41] and up to 5 days after the initial exposure [42, 43]. Much less is known, however, about the longer term transcriptional and transcriptomic gene responses to TCPOBOP and their relationship to the downstream liver pathologies that emerge, including development of CAR-dependent [44] hepatic adenomas and carcinomas following persistent TCPOBOP exposure over a 20-30 wk period [45, 46].

Here, we investigate the histopathological and transcriptomic effects of both short term and persistent CAR activation in livers from TCPOBOP-exposed mice. We show that TCPOBOP induces a dose-dependent increase in hepatic steatosis that originates pericentrally, with males more sensitive than females in the pericentral but not the periportal region, and we present a comprehensive view of both the primary and secondary gene responses and pathways that TCPOBOP dysregulates in each sex. Further, we identify upstream regulators of these gene responses, including regulators specific to TCPOBOP-exposed males, to obtain mechanistic insight into the liver pathological responses that TCPOBOP elicits in both hepatocytes and liver non-parenchymal cells. Finally, we characterize a more advanced liver pathology, including focal inflammation and immune cell infiltration, that specifically emerges when CAR is persistently activated over a 4 to 8 wk period by weekly TCPOBOP delivery using a high corn oil vehicle. TCPOBOP-treated mice may thus serve as a useful model for further investigation of the mechanisms by which foreign chemical agonists of CAR induce steatotic liver disease, as well as downstream pathologies that may emerge.

## MATERIALS AND METHODS

### Animal studies

Mouse work was conducted in accordance with ARRIVE 2.0 Essential 10 guidelines [47], including study design, sample size, randomization, experimental animals and procedures and statistical methods, and with approval from the Boston University Institutional Animal Care and Use Committee (protocol # PROTO201800698). Male and female CD-1 mice (ICR strain; strain code #022), 7-8 wk of age (average body weight: 35 g for males, 30 g for females), were purchased from Charles River Laboratories (Wilmington, MA) and housed on a 12 h light cycle (lights ON at 7:30 AM and OFF at 7:30 PM). Mice were given TCPOBOP by intraperitoneal injection at a dose ranging from 0.2 to 3 mg/kg body weight. Mice were euthanized and livers were collected at time points ranging from 1 d to 8 wk after TCPOBOP injection, as described below. Euthanasia and tissue collection were carried out between 10:30 AM and 12 noon to minimize variability due to circadian effects on liver gene expression, which impact a large subset of liver expressed genes [48–50], and is itself altered in MASH [51]. Two pieces of each liver were immediately fixed in 10% formalin; the remainder of each liver was snap frozen in liquid nitrogen for whole tissue RNA extraction (total RNA) and qPCR analysis or for extraction of liver nuclei, purification of nuclear RNA and nuclear RNA-seq analysis, as detailed below.

### TCPOBOP injection: high corn oil (vehicle) regimen

A high corn oil (vehicle) TCPOBOP dosing regimen was used in a time course study, where male livers were collected 1 wk, 2 wk, 4 wk and 8 wk after the first TCPOBOP injection (n=3 vehicle controls, and n=6 TCPOBOP-treated mice at each time point). TCPOBOP (Chem Cruze, SC-203291) was initially dissolved in 100% DMSO (Sigma, cat. # D8418) to give a stock solution at 7.5 mg TCPOBOP/ml, which was stored at -20C. Prior to use, the TCPOBOP stock solution was diluted 50-fold in corn oil to give a working solution of 0.15 mg TCPOBOP/ml of 2% DMSO/98% corn oil. Mice were injected with 20 μl of this working TCPOBOP solution per g body weight on day 0 (TCPOBOP dose: 3 mg/kg body weight), followed by additional weekly injections at one-third that dose (1 mg TCPOBOP/kg body weight) beginning on day 7, by injecting of 20 μl of 0.05 mg TCPOBOP/ml of 0.67% DMSO/99.3% corn oil. This weekly dosing schedule was designed to minimize the bioaccumulation of TCPOBOP by taking into account its 14-day half-life [52, 53]. Mice euthanized at the 8 wk time point (i.e., after a total of 8 weekly TCPOBOP injections) were found to have a depot of excess corn oil vehicle accumulating in the peritoneal cavity. An increase in liver histopathology was seen at both the 4 wk and the 8 wk TCPOBOP time points when using this regimen but was not seen in mice given weekly corn oil vehicle injections alone, or when TCPOBOP was given over the same 8 wk period but using an alternative, low corn oil vehicle regimen, described below (see Results).

### TCPOBOP injection: low corn oil (vehicle) regimen

This regimen was introduced to decrease the corn oil (vehicle) dose by > 90% as compared to the high corn oil regimen: from 20 μl corn oil/g body weight weekly to 3.6 μl (i.e., 90% of 4 μl) corn oil/g body weight every 2 wk. TCPOBOP (stock solution at 7.5 mg/ml of 100% DMSO; see above) was diluted 10-fold into corn oil to give a working solution for the low dose regimen injections. Mice were injected with 4 μl/g body weight of this solution (0.75 mg TCPOBOP/ml of 10% DMSO/90% corn oil) to deliver a TCPOBOP dose of 3 mg/kg body weight. Livers were collected 1 d, 4 d, or 2 wk later (n=4 vehicle controls, and n=6 TCPOBOP-treated mice for each sex at each time point, or as noted in individual figures). Where indicated, an additional injection of 0.375 mg TCPOBOP/ml of 10% DMSO/90% corn oil (vehicle) was given to male mice after 2, 4, and 6 wk, to achieve a bi-weekly, low corn oil regimen TCPOBOP injection dose of 1.5 mg/kg body weight. Livers were collected after 4 wk (total of 2 bi-weekly TCPOBOP injections, one on day 0, and one on day 14) or after 8 wk (total of 4 bi-weekly TCPOBOP injections, on days 0, 14, 28 and 42) (n=3 vehicle controls, and n=7 TCPOBOP-treated mice at each time point). In a separate series of experiments (dose-response study), TCPOBOP working solutions in 10% DMSO/90% corn oil were prepared at 0 (vehicle control), 0.05, 0.15 and 0.75 mg TCPOBOP/ml. Male and female mice were injected with these TCPOBOP working solutions on day 0 at 4 μl/g body weight for delivery of TCPOBOP at 0, 0.2, 0.6 and 3.0 mg/kg body weight, respectively, and livers were collected on day 14 (n=5 vehicle controls, and n=5 TCPOBOP-treated mice for each sex at each of the 4 doses).

### Tissue fixation, sectioning and staining

Freshly excised mouse liver was fixed in 10% Buffered Formalin (Fisher Scientific #23-245684) for 24 h at room temperature, transferred to 70% ethanol for 48 h then stored at 4°C until submitted to the core facility for sectioning and staining. A piece of each liver was placed in 4% formalin for fixation and subsequently stained with Hematoxylin and Eosin (H&E), Periodic Acid-Schiff with Diastase (PASD) or Sirius red, to visualize collagen networks associated with liver fibrosis. A second piece of liver was snap frozen in liquid nitrogen; tissue slices were subsequently prepared for Oil Red O staining to detect neutral lipid at the histology core of Beth Israel Deaconess Medical Center (Boston, MA) by soaking in 4% formalin, followed by soaking in 30% sucrose. A portion of each fresh liver was then prepared for cryosectioning by placement in 5 ml of 30% sucrose in PBS at room temperature until the tissue sank to the bottom. The liquid was removed by suctioning and then a thin layer of OCT was added to each cryomold (labeled with the liver ID#), and the liver placed in the center of the cryomold. OCT was carefully added, avoiding air bubbles, until it completely covered the tissue. The cryomold was then placed on a bed of dry ice for 30 min to solidify the OCT and then stored at -80°C. Following cryosectioning (5 μm slices), slides were kept at room temperature for 1 h to melt the OCT, followed by Oil Red O staining at the Beth Israel Deaconess Medical Center histology core. PAS diastase staining and trichrome staining were performed at the Experimental Pathology Laboratory Service Core (EX+) of Boston University School of Medicine. Paraffin sections were processed for antigen retrieval (Biogenex Laboratories cat. # HK0865K) then immunostained with anti-GLUL antibody (1: 1500 dilution in 3% goat serum, overnight at 4C). Sequentially cut cryosections were stained with anti-GLUL antibody and with Oil Red O, respectively, to localize the Oil Red O-stained regions of the liver lobule with respect to the GLUL-stained regions.

### Analysis of histology images

Relative hepatocyte size scores were assigned to each liver based on a set of reference images using a scale of 0 to 5 (Fig. S2B). Similarly, Oil Red O staining intensities were scored on a scale of 0 to 5 for each of n= 5 livers per group by comparing each image to a set of reference images, selected to represent the full range of staining intensities encountered in the study (Fig. S3A). The same set of reference images was used to assign scores to both periportal and pericentral regions of each liver, which allowed us to identify zone-dependent differences in both hepatocyte size and Oil Red O staining intensities for each mouse treatment group.

### Total liver cholesterol assay

A small piece of snap-frozen liver tissue (∼ 10 mg) was resuspended in 200 µL of chloroform: isopropanol: NP-40 detergent (7:11:0.1) and homogenized in an 0.4 ml glass homogenizer, then centrifuged for 5 min at 15,000 x g. The organic phase was transferred to a new Eppendorf tube, air dried at 50°C to remove chloroform, and placed in a vacuum for 30 min to remove trace organic solvent. Total cholesterol was then measured using a Cholesterol/Cholesteryl Ester Assay Kit (Abcam, cat. # ab65359).

### Blood chemistry

Blood collected from each mouse by cardiac puncture at the time of euthanasia was placed in an Eppendorf tube containing 5 mM EDTA. Plasma was obtained by centrifugation at 1500 g at 4°C for 15 min and stored at -80°C. Clinical assays for 13 analytes were performed at the Boston University Medical Center Analytical Instrumentation Core: albumin, alkaline phosphatase, alanine aminotransferase, blood urea nitrogen, calcium, cholesterol, creatinine, gamma-glutamyl transferase, glucose, phosphorus, total bilirubin, total proteins, and triglycerides.

### Isolation of liver nuclei and purification of nuclear RNA

Frozen mouse liver (∼250 mg) was placed on dry ice and minced into small pieces, which were transferred to 1 mL of Lysis buffer (100 mM Tris (pH 7.4), 146 mM NaCl, 1 mM CaCl_2_, 21 mM MgCl_2_, 0.1% NP40) in a Dounce homogenizer on wet ice. The tissue was dounced on ice (10 strokes with pestle A, then 10 strokes with pestle B) until the sample was fully homogenized. Nuclei wash buffer (1 ml of 100 mM Tris Cl (pH 7.4), 146 mM NaCl, 1 mM CaCl_2_, 21 mM MgCl_2_, 0.01% BSA, 80 U/ml Protector RNase inhibitor (Millipore Sigma cat. # 3335402001)) was added to the homogenizer, followed by gentle mixing. The sample was then passed through a 40 μm cell strainer (Sigma cat. # Z742102) sitting on top of a 50 mL centrifuge tube on ice. The homogenizer tube was rinsed with 1 ml of wash buffer, which was passed through the same 40 μm cell strainer and combined with the first strained homogenate. Each sample was kept on ice until samples from all livers were ready to proceed to the next step. Lysed cells were pelleted at 500 g for 5 min at 4°C in a swinging bucket rotor. The pellets containing crude nuclei were resuspended in 1000 μl Staining Buffer (PBS, 2% BSA (Sigma cat. # SRE0036), 80 U/ml Protector RNase Inhibitor). Samples were centrifuged at 500 g for 5 min at 4°C in a swinging bucket rotor. The pelleted nuclei were resuspended in 1000 μl of Staining Buffer and passed through a 20 μm filter (PluriSelect cat. # 43-50020-50) then centrifuged at 500 g for 5 min at 4°C in a swinging bucket rotor. The purified nuclei were resuspended in 250 μl nuclease-free water, followed by the immediate addition of 750 μl Trizol-LS, pipetting up and down thoroughly to mix, followed by RNA isolation using the manufacturer’s protocol (Life Technologies).

### RNA-seq and differential expression analysis

RNA-seq analysis was performed using nuclear RNA extracted from frozen livers from n=3-4 individual mice per treatment group. Sequencing libraries were prepared starting with 1 μg of liver nuclear RNA, by poly(A) selection using the NEBNext Poly(A) mRNA Magnetic Isolation Module, followed by processing with the NEBNext Ultra Directional RNA Sequencing for Illumina kit (New England Biolabs). Illumina sequencing, 150 paired end reads, was performed at Novogene Corporation Inc. to a mean depth of 21.5 million read pairs per RNA-seq library (Table S1A). Data were analyzed using a custom RNA-seq analysis pipeline, including TopHat for mapping sequence reads to the mouse genome (release mm9), featureCounts to obtain read counts for 24,197 RefSeq genes, and edgeR to identify differentially expressed genes at FDR < 0.05 [40]. Raw Fastq files and processed data files are listed in Table S1A and are available at GEO (https://www.ncbi.nlm.nih.gov/geo/), accession # GSE248858. Full datasets for differentially expressed genes are provided in Table S1B and Table S4A.

### Ingenuity Pathway Analysis (IPA)

Genes showing significant differential expression in each sex and at each time point of TCPOBOP exposure were submitted to IPA (Qiagen, Inc) to identify enriched canonical pathways (Table S2), enriched upstream regulators (Table S3) and enriched Disease and Bio Functions and Tox Functions (Table S5). These analyses gave p-values (Benjamini-Hochberg corrected, where indicated), which indicate the probability of association of the input genes with the pathway, upstream regulator or other function by random chance alone, as well as a Z-score, whose directionality indicates the activation state or the inhibition state of the pathway or the upstream regulator. Terms with |Z-score| >2 are considered significant; terms with |Z-score| ≤ 2 are of indeterminant directionality. Upstream regulators with a molecular type identified as chemical or biological were excluded from all downstream analysis.

### DAVID analysis

Functional enrichment analysis of differentially expressed gene sets was performed using DAVID (https://david.ncifcrf.gov/tools.jsp) [54] with default parameters, except that Gene Ontology (GO) FAT terms were used in place of GO DIRECT terms to include a broader range of enrichment terms, which are excluded by the default GO DIRECT option.

### qPCR analysis

Total liver RNA was purified from ∼100 mg of frozen liver tissue using TRIZOL, following the manufacturer’s protocol (Life Technologies). cDNA synthesis was performed using cDNA Reverse Transcription Kit (Fisher, cat. #43-688-14) with 1 μg of purified total liver RNA. Quantitative real time PCR (qPCR) was performed using Power SYBR Green PCR Master Mix (ThermoFisher) in 384-well plates and assayed using an CFX384 Touch Real-Time PCR Detection System (Bio-Rad) and gene specific primers shown in Table S1E. Fold-change values were calculated using the ΔΔCt method. The expression of 18S ribosomal RNA (Ct value) was used to normalize liver cDNA samples.

### Statistics

Significance was assessed by ANOVA with Tukey’s multiple comparisons, or multiple t-test-corrected p-values implemented in GraphPad Prism using the Sidak method, with alpha = 0.05, and without assumption of a consistent standard deviation between groups, as indicated in each figure. Data are presented as the mean ± SEM values based on biological replicate livers. The significance of enrichment of TCPOBOP-responsive gene sets for binding of CAR, which identifies primary TCPOBOP-responsive CAR target genes, was computed using Fisher Exact test, which was applied to ChIP-seq data for CAR binding to mouse liver chromatin reported previously [29, 30].

## Results

### TCPOBOP induces dose-dependent increases in liver index and hepatocyte cell size

Male mice given a single, receptor-saturating dose of TCPOBOP (3 mg/kg body weight, i.p.) showed an increase in liver/body weight ratio (liver index) after 4 d (Fig. 1A). This increase was maximal by 1-2 wk, with no further increases seen when mice were given additional, weekly TCPOBOP injections and then examined after 4 or 8 wk. In male mice, liver index increased significantly 2 wk after a single injection of TCPOBOP at a 0.2 mg/kg dose, corresponding to the ED50 for transcriptional activation of *Cyp2b10* [53], whereas in female mice, a significant increase in liver index was first seen at 0.6 mg TCPOBOP/kg (Fig. S1).

**Fig. 1.**
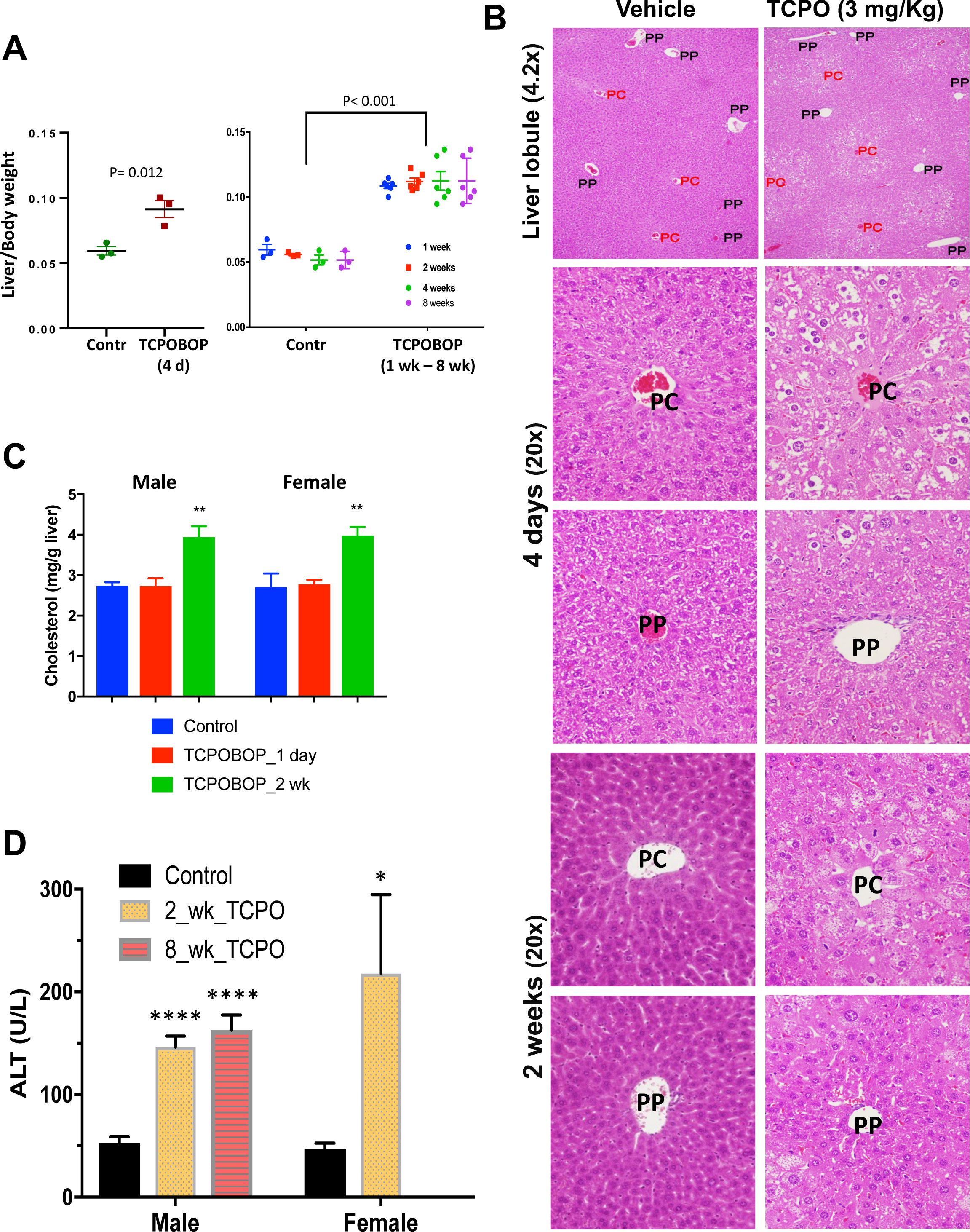
Liver pathology induced by prolonged TCPOBOP exposure. **(A)** Liver to body weight ratio (liver index) of mice treated with TCPOBOP for times ranging from 4 d to 8 wk, with TCPBOP given using a low corn oil vehicle regimen (4 d time point) or weekly injections using a high corn oil vehicle regimen (1-8 wk). Each data point represents an individual mouse. **(B)** H&E staining of liver sections 4 d or 2 wk after a single TCPOBOP injection (3 mg/kg, low corn oil regimen), with liver lobule pericentral (PC) and periportal (PP) regions marked. Enlarged hepatocytes are specific to the pericentral region. **(C)** TCPOBOP (low corn oil regimen) increases hepatic total cholesterol content after 2 wk in both sexes (p< 0.005, 2-way ANOVA, with Tukey’s multiple comparisons test) compared to male or female controls, mean ± SEM (n=3-4 livers/group). **(D)** TCPOBOP (low corn oil regimen) increases plasma ALT (alanine aminotransferase) levels, mean ± SEM (n=4-10 livers/group). Shown are multiple t-test-corrected p-values compared to sex-matched controls: ****, p < 0.0001; *, p < 0.05. One TCPOBOP-treated female had a plasma ALT value > 1,000 and was omitted from the analysis.

Hepatocyte cell size increased in a liver lobule zone-dependent manner, as seen 4 d after a single TCPOBOP injection. The effect persisted 2 wk after a single TCPOBOP injection, with hepatocyte hypertrophy characterizing cells near the central vein (pericentral hepatocytes), the liver lobule zone where CAR, the nuclear receptor for TCPOBOP, shows highest expression [41], but was much less apparent near the portal triad (periportal hepatocytes) (Fig. 1B, Fig. S2). Moreover, pericentral hepatocyte hypertrophy was greater in male than female liver (Fig. S2B). Nuclei of TCPOBOP-exposed hepatocytes retained their characteristic round shape; they did not display the hepatocyte nuclear membrane deformation reported for a dietary model of mouse MASLD and in hepatocytes from MASLD patients, which was proposed to contribute to the activation of repressed genomic regions containing lipogenic genes in steatotic liver disease [55].

### Sex-dependent zonation of TCPOBOP-induced steatosis

Neutral lipid accumulation in hepatocytes, and the pathogenesis of MASLD, is accompanied by dysregulation of hepatic cholesterol homeostasis and liver cholesterol accumulation [56]. Consistent with this, liver cholesterol content was significantly increased in livers of both male and female mice 2 wk after a single TCPOBOP injection (Fig. 1C) and remained elevated after 8 weekly injections (Fig. S1C). Oil Red O staining revealed a striking, dose-dependent accumulation of neutral lipid in pericentral region hepatocytes, as was seen 2 wk after TCPOBOP injection (Fig. 2, Fig. S3A). pericentral hepatocyte lipid accumulation began within 4 d of TCPOBOP injection, as indicated by the whitish regions surrounding each central vein in H&E-stained images (Fig. 1B). Moreover, clear sex differences in the patterns of TCPOBOP-induced lipid accumulation were apparent, with Oil Red O staining in the pericentral region lower in female than in male liver. In contrast, in the periportal region, Oil Red O staining intensity was greater in female liver (Fig. 2, Fig. S3A; sex differences in both regions significant by 2-way ANOVA at p<0.001, Fig. S3B). Accordingly, male but not female livers showed greater lipid accumulation in pericentral than in periportal hepatocytes at all three TCPOBOP doses (Fig. S3C). Furthermore, periportal distal male hepatocytes showed greater lipid accumulation than periportal proximal cells, at both 2 wk and 8 wk after initiating TCPOBOP treatment (Fig. S3D). These zone-dependent sex-differences in the steatotic effects of TCPOBOP are consistent with, and may be explained by, sex differences in the zonation of CAR expression revealed by single cell-based RNA-seq [41]. Specifically, in male mouse liver, CAR showed a significant, 4.3-fold higher expression in pericentral than in periportal hepatocytes (p = 3.9E-12), whereas in female liver the zonation bias in CAR expression (pericentral > periportal, < 2-fold) did not reach significance due to a 2.2-fold higher basal level of CAR expression in periportal hepatocytes in female compared to male liver (Fig. S4). The latter finding is consistent with the greater susceptibility of female periportal hepatocytes to TCPOBOP-induced lipid accumulation seen in Fig. 2. These sex differences in the zonation of lipid accumulation, most notably the elevated lipid levels in periportal hepatocytes from TCPOBOP-treated female mice, were also evident from the checkered appearance of the overall liver lobule pattern of Oil Red O staining seen in 4.2x images of male liver but not female liver (Fig. 2, top 2 rows).

**Fig. 2.**
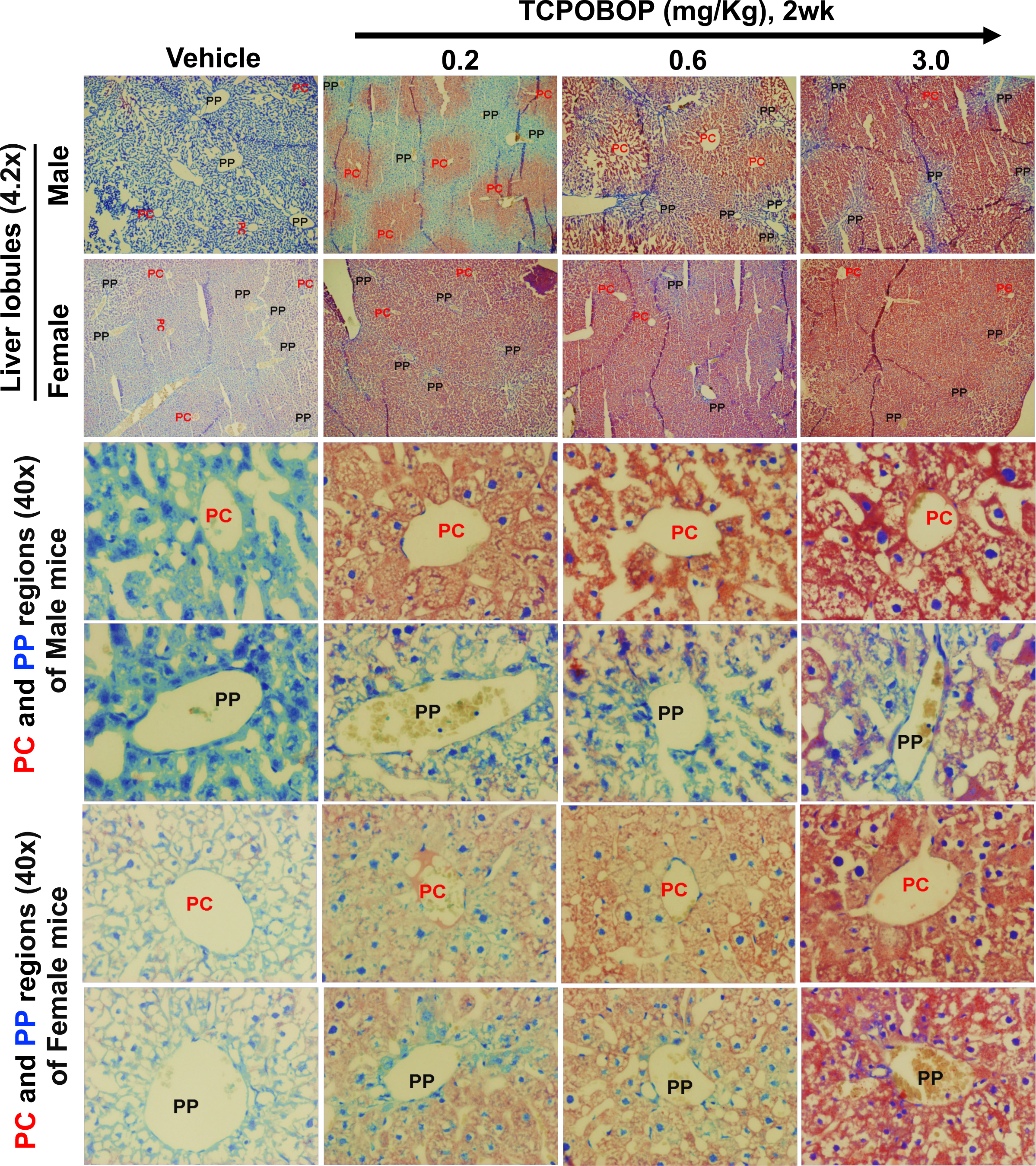
Neutral lipid staining of TCPOBOP-exposed livers from male and female mice. Mice were given a single injection of TCPOBOP at the indicated doses, or vehicle control (low corn oil regimen) and euthanized 2 wk later. Liver sections were stained with Oil Red O. Representative images (based on analysis of all 5 livers per treatment group) are shown for both the periportal (PP) and pericentral (PC) regions, as marked. See Fig. S3A-S3C for quantification of staining intensities.

We validated the zonated increase in neutral lipids using the pericentral marker protein glutamine synthetase (*Glul*, glutamine ammonia lyase), which detoxifies ammonia entering hepatic circulation and is selectively expressed in the first 1-3 layers of perivenular hepatocytes surrounding the central vein in untreated liver [57, 58]. First, we stained paraffin-embedded sections to identify GLUL-positive pericentral hepatocytes and discovered that 2 wk TCPOBOP exposure disrupts the cellular profile of GLUL-positive staining, which became more diffuse and less highly localized to the immediate vicinity of the central vein (Fig. S5A, *left*; Fig. S5B). Since lipids are substantially extracted during the preparation of paraffin-embedded tissue blocks, we used liver cryosections to examine two sequential slices from the same liver. One section was stained with Oil Red O, and the next section was stained with anti-GLUL antibody (Fig. S5A, *middle* and *right*). The GLUL-positive distribution pattern matched the pattern of Oil Red O staining, both in vehicle control liver (basal Oil Red O staining) and in TCPOBOP-treated liver (Fig. S5A, *top* and *bottom* rows), verifying that TCPOBOP-stimulated lipid accumulation is largely localizes to, and likely originates in the pericentral zone.

Thus, male mice are more susceptible than female mice to the effects of TCPOBOP on liver index, pericentral hepatocyte size and pericentral neutral lipid accumulation (hepatosteatosis), whereas female mice show increased lipid accumulation in the periportal region compared to males. In both sexes, TCPOBOP increased circulating levels of alanine aminotransferase, which is indicative of hepatocyte damage, beginning 2 wk after TCPOBOP treatment (Fig. 1D). However, we did not detect any increase in liver fibrosis, as judged by trichrome and Sirius red staining 2 wk or 4 wk after initiating TCPOBOP injection (data not shown). Circulating levels of 12 other plasma analytes (see Methods) were unchanged by TCPOBOP exposure.

### TCPOBOP induces MASLD-associated genes

We used qPCR to investigate the impact of TCPOBOP exposure on select genes. *Elovl6*, which plays a role in the elongation of C12-C16 saturated and monounsaturated fatty acids, was significantly increased in expression in male liver, both 1 d and 2 wk after TCPOBOP injection; females showed the same trends but did not reach statistical significance. Liver pyruvate kinase liver/red blood cell (*Pklr*), a MASLD driver gene that promotes steatosis and liver fibrosis [59, 60], showed increased expression 2 wk after TCPOBOP injection in both sexes. *Pnpla3*, which is expressed in a female-biased manner and whose common genetic variant (I148M) is a major contributor to inherited MASLD susceptibility in women [61–63], showed a trend of increased expression in females after 2 wk TCPOBOP exposure. These responses can be compared to the effects of TCPOBOP on *lnc13509* [40], whose hepatic RNA level increased up to 40-fold within 1 d of TCPOBOP exposure and persisted at 2 wk (Fig. 3A). Other genes associated with liver pathology that were induced by TCPOBOP after 2 wk, but not after 1 d exposure, included *Gpnmb*, *Mmp12* and *Col1a1* (Fig. 3B, and data not shown). Gpnmb is a macrophage-specific transmembrane glycoprotein that negatively regulates inflammation [64], and the matrix metalloproteinase Mmp12 and the major hepatic collagen subtype Col1a1 have both been implicated in regulation of inflammation and hepatic fibrosis [65, 66]. All 3 genes are up regulated in a diet-induced animal model of MASLD [67].

**Fig. 3.**
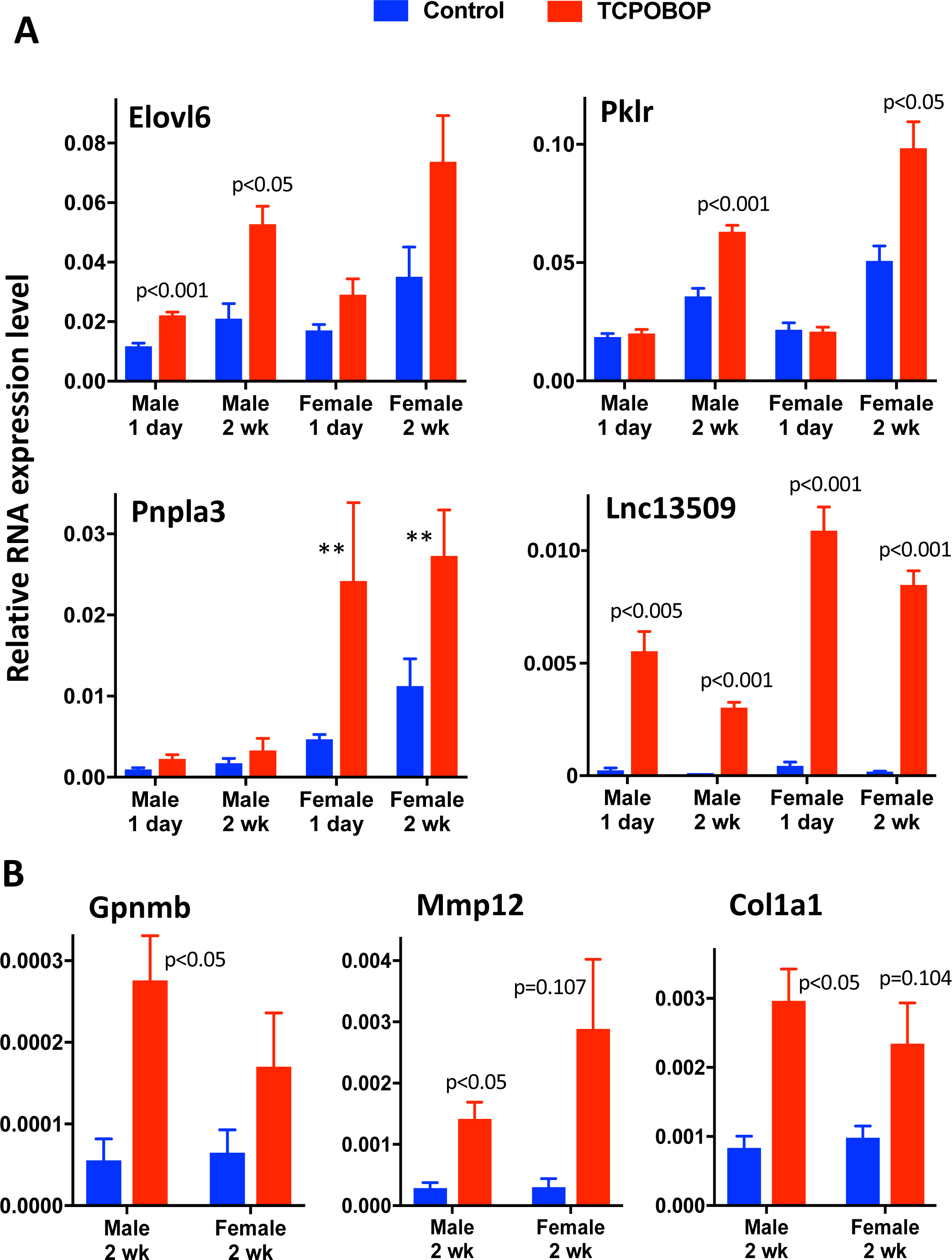
qPCR analysis of TCPOBOP-responsive RNAs with diverse functions related to hepatic lipid metabolism, inflammation and fibrosis. Data shown are relative RNA levels after 1 d or 2 wk TCPOBOP exposure (low corn oil regimen), mean ± SEM values for n=4 livers/group (vehicle controls) or n=6 livers/group (TCPOBOP-treated livers) **(A)**. Statistical significance values shown are multiple t-test-corrected p-values for TCPOBOP vs vehicle control, using the Holm-Sidak method; unmarked TCPOBOP vs control comparisons were not significant. **, sex difference significant at p < 0.01 for TCPOBOP-exposed livers by 2-way ANOVA. **(B)** Analysis as in (A). No significant TCPOBOP-induced changes in expression were seen at 1 d for Gpnmb, Mmp12 or Col1a1 (not shown).

### Global transcriptional responses to CAR activation

Nuclear RNA-seq analysis was carried out to characterize TCPOBOP-induced transcriptional responses globally, for both male and female mouse livers collected either 1 d or 2 wk after a single TCPOBOP injection (low corn oil regimen). Several hundred RefSeq genes were dysregulated by TCPOBOP after 1 d, consistent with our prior findings [40]. More genes responded to TCPOBOP in female than in male liver at both time points (Fig. 4A, Table S1B). A majority of genes that responded in common in both sexes were more strongly induced, or were more strongly repressed, in female as compared to male liver (Fig. S6), perhaps due to the higher overall level of CAR in female liver [41, 68, 69]. In male, but not female liver, more genes were up regulated than were down regulated by TCPOBOP (1 d TCPOBOP: up/down regulation gene ratio = 1.94 in male liver vs. 1.13 in female liver; 2 wk TCPOBOP: up/down regulation gene ratio = 1.42 in male vs. 1.09 in female liver).

**Fig. 4.**
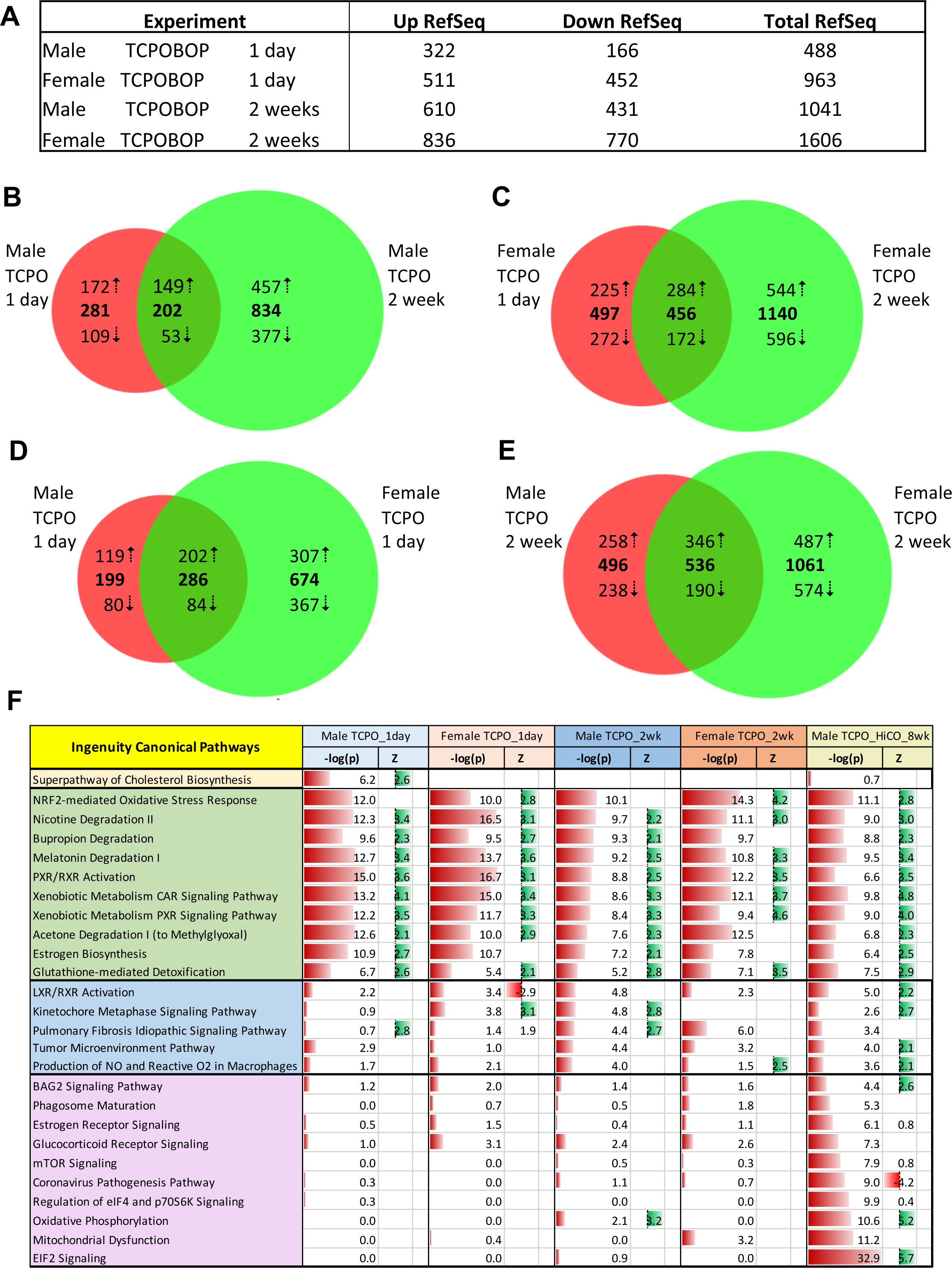
Nuclear RNA-seq identifies RefSeq genes responsive to TCPOBOP. **(A)** Numbers of RefSeq genes that responded to 1 d or 2 wk TCPOBOP treatment (3 mg/kg, low corn oil regimen) in male or female mouse liver at FDR < 0.05. **(B-E)** Venn diagrams showing numbers of TCPOBOP-responsive genes that overlap between the two time points (B, C) in each sex, or between males and females (D, E) at each time point. Bold, total number of up + down regulated genes in each Venn diagram segment (*up* and *down* arrows, respectively). (A), but not (B-E) includes small numbers of genes that show a discrepancy in the direction of TCPOBOP response between time points or between sexes (5, 10, 3 and 9 genes for panels B-E, respectively). For example, 5 of 207 genes responding to TCPOBOP in male liver at both time points show opposite responses at 1 d vs 2 wk and are excluded from (B). See Table S1B for full gene listings. (**F)** Representative IPA canonical pathways enriched for at least 1 of the 5 indicated sets of TCPOBOP-responsive genes at -log10 p-value ≥ 4. Pathways are organized into 4 groups, namely, those enriched for genes responsive to TCPOBOP after 1 d only (top row, yellow), those enriched for genes responsive in all 5 conditions (green), those enriched for genes responsive at 2 wk in male liver, but not at 1 d (blue), and those enriched for genes responsive only at 8 wk when using the high corn oil (HiCO) regimen (purple). Z-scores > 2 (short green bars, to the right) indicate up regulation of the pathway and Z-scores < 2 indicate down regulation of the pathway (short red bars, to the left). Full dataset is shown in Table S2.

We observed a large increase in the number of TCPOBOP-responsive genes between 1 d and 2 wk (Fig. 4A-4C, Table S1B). Overall, 113 genes were consistently up regulated by TCPOBOP in both sexes at both time points, while 27 genes were consistently down regulated (Table S1B, column L). Gene inductions ranged up to 1,000-fold and included well characterized genes active in drug and xenobiotic metabolism (e.g., *Cyp2b10*, *Cyp2c55*, *Gstm3m3*, *Akr1b7*). Novel TCPOBOP-induced genes of special interest include *Pnliprp1* and *Fzd10*, whose expression increased >90-100-fold in both sexes, both 1 d and 2 wk after a single TCPOBOP injection. Pnliprp1 is a metabolic inhibitor of triglyceride digestion whose expression is positively associated with diets rich in high fat intake across species [70, 71]. Fzd10 is a plasma membrane receptor whose signaling activates beta-catenin and Yap1. Fzd10 promotes tumorigenicity and metastasis of liver cancer stem cells [72] and could be a key early factor in the hepatocarcinogenesis seen after prolonged TCPOBOP exposure [46]. Importantly, 34 of the 113 consistently early/persistently up regulated genes are Lipid Metabolic Process genes (GO:000662934; enrichment Benjamini-corrected p = 2.1E-10 by DAVID analysis), which may contribute to the observed hepatosteatosis. A total of 138 lipid metabolic process genes responded to TCPOBOP after either 1 d or 2 wk; these genes constitutes the most highly enriched DAVID cluster at both time points in both sexes (enrichment scores = 10.9-23-fold) (Table S1A, column M).

### Genes showing delayed response to TCPOBOP are enriched for indirect CAR genes targets with MASLD-related functions

TCPOBOP rapidly activates the transcription of many genes in mouse liver, as shown by RNA-seq analysis of nuclear RNA extracted 3 h after a single i.p. injection of TCPOBOP at 3 mg/kg [40]. The direct nature of these gene responses is supported by the rapid chromatin opening that occurs at their nearby enhancer sequences, many of which harbor binding sites for TCPOBOP-activated CAR [29]. Similarly, the 1 d TCPOBOP-responsive genes identified in the present study were highly enriched for direct CAR binding, with enrichment scores (ES) = 6.2-fold (p< 1E-05, Fisher Exact test) and 4.5-fold (p< 1E-05) for 1 d TCPOBOP responses in male and female liver, respectively, when compared to a background gene set comprised of genes stringently unresponsive to TCPOBOP treatment Table S1C). The enrichment for CAR binding increased to ES = 9.0-fold (p< 1E-05) when genes that responded to 1 d TCPOBOP in both sexes were considered (Table S1C). In contrast, the set of genes induced by TCPOBOP after 2 wk but not after 1 d exposure (late response genes) showed either no significant enrichment for CAR binding (genes unresponsive to 2 wk TCPOBOP in males, or genes unresponsive in both sexes; p > 0.05) or weak enrichment (genes unresponsive to 2 wk TCPOBOP in females, ES = 1.98, p = 0.028) (Table S1C). We conclude that many of the late TCPOBOP-responsive genes are not induced by a direct CAR binding mechanism, i.e., they are indirect, secondary response genes.

151 TCPOBOP-induced genes were identified as late response genes in livers of both male and female mice (Table S1B, column L). These genes were significantly enriched for response to cytokine (FDR = 2.1E-03) and innate immune response (FDR = 7.3E-03), among others (DAVID analysis, Table S1D). This finding is reminiscent of the secondary activation of inflammation and immune response pathways in MASLD [73, 74]. Many of these late responding/secondary response genes have biological activities related to liver steatosis and other, downstream pathologies, including those involving liver non-parenchymal cells. One such gene is *Bmp8b* (9-15-fold induction by TCPOBOP at 2 wk), which is up regulated in livers of mice fed a Western diet and promotes a hepatic stellate cell proinflammatory phenotype that contributes to the progression of non-alcoholic steatohepatitis [75]. Another late response gene, *Ubd* (5-14-fold induction at 2 wk), is a ubiquitin-like protein that is up regulated in patients with MASLD [76] and is over expressed in 70% of human HCC patients [77]. Other TCPOBOP late response genes may ameliorate the severity of liver pathology; examples include *Arg2* (4-8-fold induction), which can suppress spontaneous steatohepatitis [78], and *Ppp1r3g* (6-14-fold induction), whose overexpression abrogates alcohol-induced hepatic lipid deposition [79]. Late response genes noted above (Fig. 3B) include the MASH-associated macrophage markers *Gpnmb* and *Mmp12* [11, 80], and the profibrogenic marker for activated hepatic stellate cells, *Col1a1* [81].

### Canonical pathway analysis and Disease, Bio and Tox Function analysis

The full set of TCPOBOP responsive genes (FDR< 0.05) was analyzed using IPA software to identify significantly enriched canonical pathways, almost all showing up regulation (positive Z-scores; Fig. 4F, Table S2). Pathways related to CAR, PXR and xenobiotic metabolism dominated the list of pathways enriched at both 1 d and 2 wk in both sexes (Fig. 4F, *green*). Pathways associated with protection from oxidative stress, including NRF2-mediated oxidative stress response and glutathione-mediated detoxification, were also highly enriched. Pathways that were more significantly enriched in male liver after 2 wk TCPOBOP exposure as compared to 1 d exposure included LXR/RXR activation, which promotes lipogenesis [82], and pulmonary fibrosis, tumor microenvironment, and production of nitric oxide and reactive oxygen in macrophages, which may contribute to liver damage (Fig. 4F, *blue*). Consistent with this, IPA Disease and Bio Function analysis revealed that macrophage activation and immune response/immune-mediated inflammatory disease were highly induced after 2 wk but not after 1 d TCPOBOP exposure, as was reactive oxygen species production and metabolism (Table S5A). Top enriched Tox Function categories identified at both TCPOBOP time points included liver hyperplasia and liver steatosis, both consistent with the observed histopathology, and HCC; whereas the Tox Functions liver damage and liver inflammation showed much stronger enrichment at 2 wk than at 1 d in both sexes (Table S5B). Interestingly, liver fibrosis was exclusively associated with 2 wk TCPOBOP exposure in both sexes (Table S5B), despite the absence of detectable fibrosis when assayed by Sirius red staining, as noted above.

### Early upstream regulators include transcription factors and other responders to liver injury and inflammation

IPA analysis identified both activated (Z-score > 2) and inhibited (Z-score < -2) upstream regulators, which are predicted to control the gene response pathways activated by TCPOBOP (Fig. 5, Table S3). The most highly significant activated upstream regulators across both sexes at both time points (-log10 (B-H p-value) > 4; Fig. 5A) included CAR itself (Nr1i3) and the closely related PXR (Nr1i2), as expected, as well as the liver transcription factor CEBPB, whose binding is strongly enriched at liver chromatin regions that open following TCPOBOP exposure [29]. Another early activated upstream regulator, NFE2L2 (NRF2), was previously linked to CAR activation and induces antioxidant response element-containing genes involved in injury and inflammatory responses [83]. Other early activated upstream regulators reported previously for 1 d TCPOBOP-exposed livers [40] include the inflammatory damage markers TNF, IL6, and LEP, as well as CTNNB1 (β-catenin), which contributes to CAR-induced hepatocyte proliferation [84], and NFKBIA, a target of the transcriptional factor NFκB, which regulates many inflammatory and immune responses. Early upstream regulators whose activity was consistently inhibited by TCPOBOP include the anti-oxidant enzymes GSR (glutathione reductase) and TXNRD1 (thioredoxin reductase 1), indicating decreased protection from MASLD progression-associated by oxidative stress. Two NRs were identified as early inhibited upstream regulators, NR1H4 (FXR) and NR0B2 (small heterodimer partner). Combined deletion of these two NR genes leads to activation of CAR as well as intrahepatic cholestasis [85]. The inverse relationship between CAR activity and that of these two NRs may in part be due to their competition for CAR binding sites in liver chromatin, which was shown experimentally for NR1H4 [31]. Early upstream regulators that show an inconsistent activation status in TCPOBOP-exposed liver (2 > Z-score > -2) include the transcription factor Ahr, six other NRs (ESR1, PPARA, PPARG, RORA, RORC, HNF4), and the growth hormone-activated transcription factor STAT5B. STAT5B is a global regulator of genes showing sex-biased expression in mouse liver [86, 87], many of which are dysregulated by TCPOBOP in a sex-dependent manner [40].

**Fig. 5.**
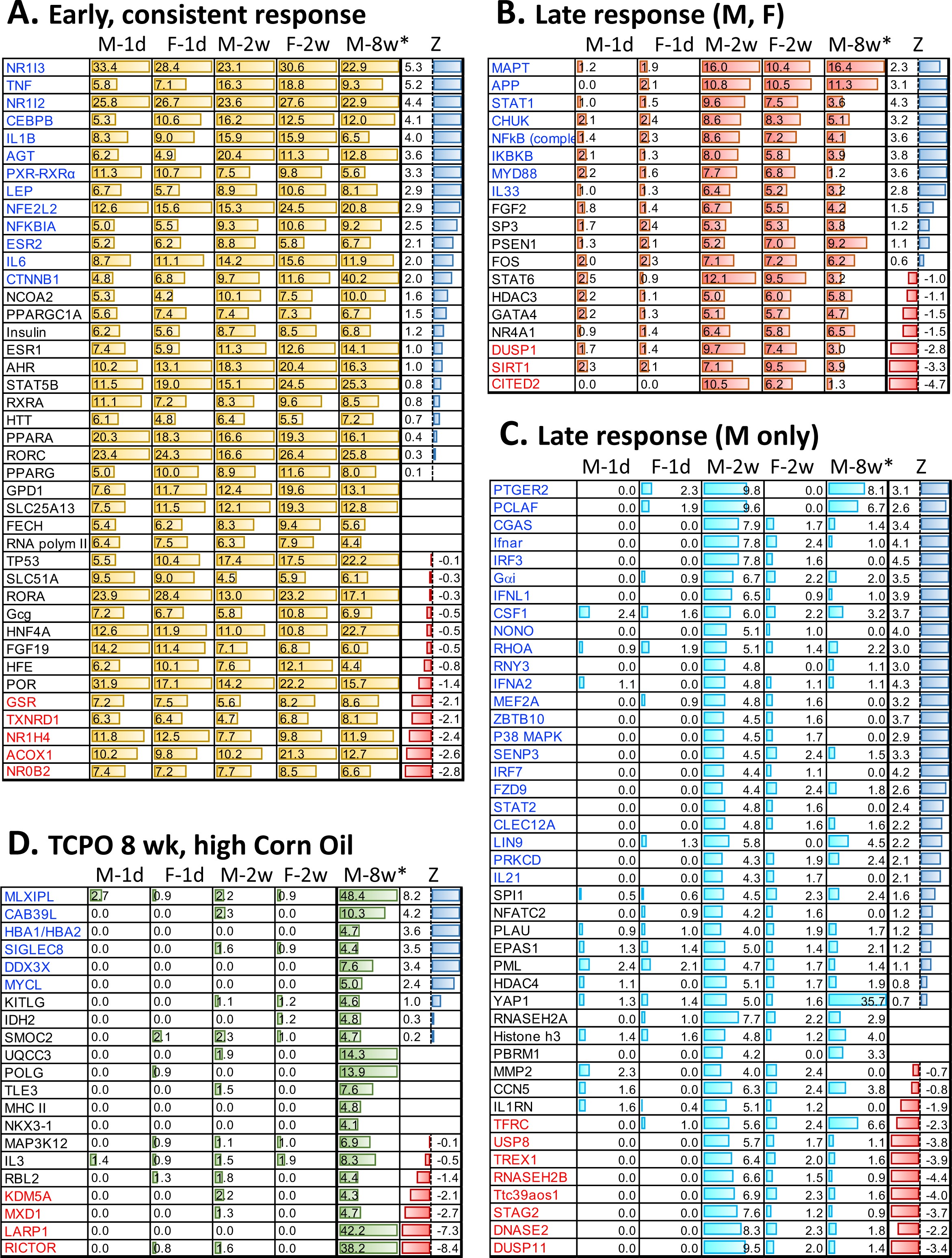
Upstream regulators of TCPOBOP-responsive genes that are significant, as determined by IPA Upstream Regulator analysis, and that met these criteria: **(A)** -log10 (Benjamini-Hochberg-corrected (B-H) p-value) > 4 for all 5 datasets (as shown above each column), i.e., are consistent across TCPOBOP gene responses in both sexes and at all time points examined; **(B)** -log10 (B-H) p-value > 5 for both male and female 2 wk TCPOBOP datasets and -log10 (B-H p-value) < 2.5 for both male and female 1 d TCPOBOP datasets, i.e., upstream regulators specific for late (2 wk) TCPOBOP-responding genes; **(C)** -log10 (B-H p-value) > 4 for the male 2 wk TCPOBOP dataset and -log10 (B-H p-value) < 2.5 for the female 2 wk and for both the male and female 1 d TCPOBOP datasets, i.e., upstream regulators specific for the late (2 wk) TCPOBOP-responding genes in male but not female liver; or **(D)** -log10 (B-H p-value) > 4 for the male 8 wk TCPOBOP/high corn oil regimen dataset (*) and -log10 (B-H p-value) < 2.5 for the 4 other data sets (exception: MLXIPL value = 2.7 in male 1 d dataset), i.e., upstream regulators specific for the male 8 wk TCPOBOP/high corn oil regimen. Z-scores shown are for the male 2 wk TCPOBOP dataset, except for (D), where the values shown are for the male 8 wk TCPOBOP/high corn oil dataset. Z > 2 indicates up regulation of the upstream regulator (gene names in *blue*) and Z < 2 indicates down regulation (gene names in *red*). Of the 21 upstream regulators shown in (D), only KITLG and SMOC2 met the criteria of -log10 p-value of overlap >4 in male 8 wk TCPOBOP/low corn oil regimen livers. Full dataset is shown in Table S3.

### Late-responding upstream regulators are associated with inflammation and MASLD progression

To better understand the expansion of liver transcriptional responses to TCPOBOP over time (Fig. 4B, Fig. 4C), we identified robust late upstream regulators as those that were highly enriched in 2 wk TCPOBOP liver (-log10 (B-H p-value) > 5) in both sexes but displayed weak or no enrichment in 1 d TCPOBOP livers of both sexes (-log10 (B-H p-value) < 2.5) (Fig. 5B). Consistent with our finding, above, that macrophage activation and immune response/inflammatory disease Bio Functions were specifically induced after 2 wk TCPOBOP exposure (Table S5A), several regulators of immune response and cytokine production were identified as activated late upstream regulators (Fig. 5B). These include STAT1, which promotes MASH [88, 89], MYD88, whose persistent activation induces liver inflammation and M2 macrophage polarization, promoting HCC [90, 91], the cytokine IL33, which has both pro- and anti-inflammatory properties and is released in response to cell damage and necrosis [92], and the inflammatory pathway regulators NFKB, IKBKB, and CHUK [93]. Inhibited late upstream regulators include two genes whose inhibition may contribute to steatotic liver disease and its progression. One gene, DUSP1, is a major negative regulator of MAP kinase signaling that is decreased in MASLD patients and shows increased expression following gastrectomy linked to the amelioration of liver disease [94]. The second gene, SIRT1, protects cells from metabolic stress and steatotic liver disease by deacetylating proteins associated with lipid metabolism [95, 96]. A third inhibited late upstream regulator, CITED2, is a transcriptional co-activator that promotes hepatic gluconeogenesis [97].

### Late-responding upstream regulators specific to male liver give mechanistic insight into male-biased liver pathology

Given the greater hypertrophic and steatotic responses seen in 2 wk TCPOBOP-exposed male compared to female liver, we investigated upstream regulators specifically associated with 2 wk TCPOBOP male liver. We identified 44 such regulators, of which 23 were predicted to be activated and 8 were inhibited (Fig. 5C). Remarkably, the full set of 44 regulators showed significant enrichment for specific top GO terms, including regulation of cell proliferation (FDR = 1.27 E-04), DNA metabolic process (FDR = 3.28E-05), defense response (FDR = 8.60E-05), chromatin binding (FDR 9.75E-06), and cellular response to cytokine stimulus (FDR = 3.28E-05) (Table S3D). Activated upstream regulators of interest specific to 2 wk TCPOBOP-exposed male liver include multiple factors specifically linked to either MASLD or HCC: CGAS, which facilitates type-I interferon production via the STING pathway [98] and is activated in MASLD by replication stress [99]; SENP3, whose increased expression in MASLD is positively associated with hepatocyte lipid accumulation [100]; PTGER2, which is up regulated in HCC [101]; RHOA, which promotes tumor cell proliferation and metastasis and is a poor prognostic factor for HCC [102]; NONO, a scaffold protein that binds Neat1, sequesters other RNAs in paraspeckles [103] and contributes to HCC progression [104]; and the protein kinase C gene PRKCD, which is associated with poor overall survival in HCC [105]. Strikingly, 5 of the 8 upstream regulators whose functions are specifically inhibited in 2 wk TCPOBOP-exposed male but not female liver have anti-inflammatory activity, namely: the de-ubiquitination enzyme USP8, whose decreased activity is associated with increased liver macrophage (Kupffer cell) inflammation and increased liver fibrosis [106], but whose inhibition is therapeutically beneficial in the context of HCC [107, 108]; the nucleases TREX1 and DNASE2, whose down regulation leads to cytoplasmic accumulation of nuclear DNA, CGAS-STING activation, and HCC promotion in a high fat dietary mouse model [109]; Ttc39aos1, an anti-inflammatory lncRNA that represses transcription of immune response genes [110]; and the RNA phosphatase DUSP11, whose deficiency leads to significantly increased production of inflammatory cytokines following lipopolysaccharide treatment [111]. The inhibition of these upstream regulators in 2 wk TCPOBOP-exposed male liver provides mechanistic insight into the pathological responses to TCPOBOP activation, most notably inflammatory responses that are predicted to be preferentially activated in male liver.

### Combination TCPOBOP + high corn oil exposure increases liver pathology and dysregulates unique upstream regulators

In a separate set of experiments, male mice were given TCPOBOP on a weekly dosing schedule delivered using a high corn oil vehicle regimen (20 μl corn oil/g body weight per week). Using this regimen, we observed accumulation of corn oil in the peritoneal cavity after 8 wk, as well as more advanced TCPOBOP-induced liver pathology when compared to the standard low corn oil vehicle TCPOBOP regimen (i.e., 3.6 μl corn oil/g body weight every 2 wk, corresponding to a 90% lower total dose over an 8 wk exposure period). Specifically, weekly TCPOBOP treatment using the high corn oil regimen induced focal inflammation and immune cell infiltration, beginning at 2 wk and continuing at the 4 wk and 8 wk time points. TCPOBOP also induced a time-dependent increase in liver sinusoidal space in the high corn oil group that was readily evident after 8 wk (Fig. 6, Fig. S7). This increase in sinusoidal space may alter hepatic blood flow or permeability and is consistent with more advanced pathology seen in some but not all models of liver disease [112–115].

**Fig. 6.**
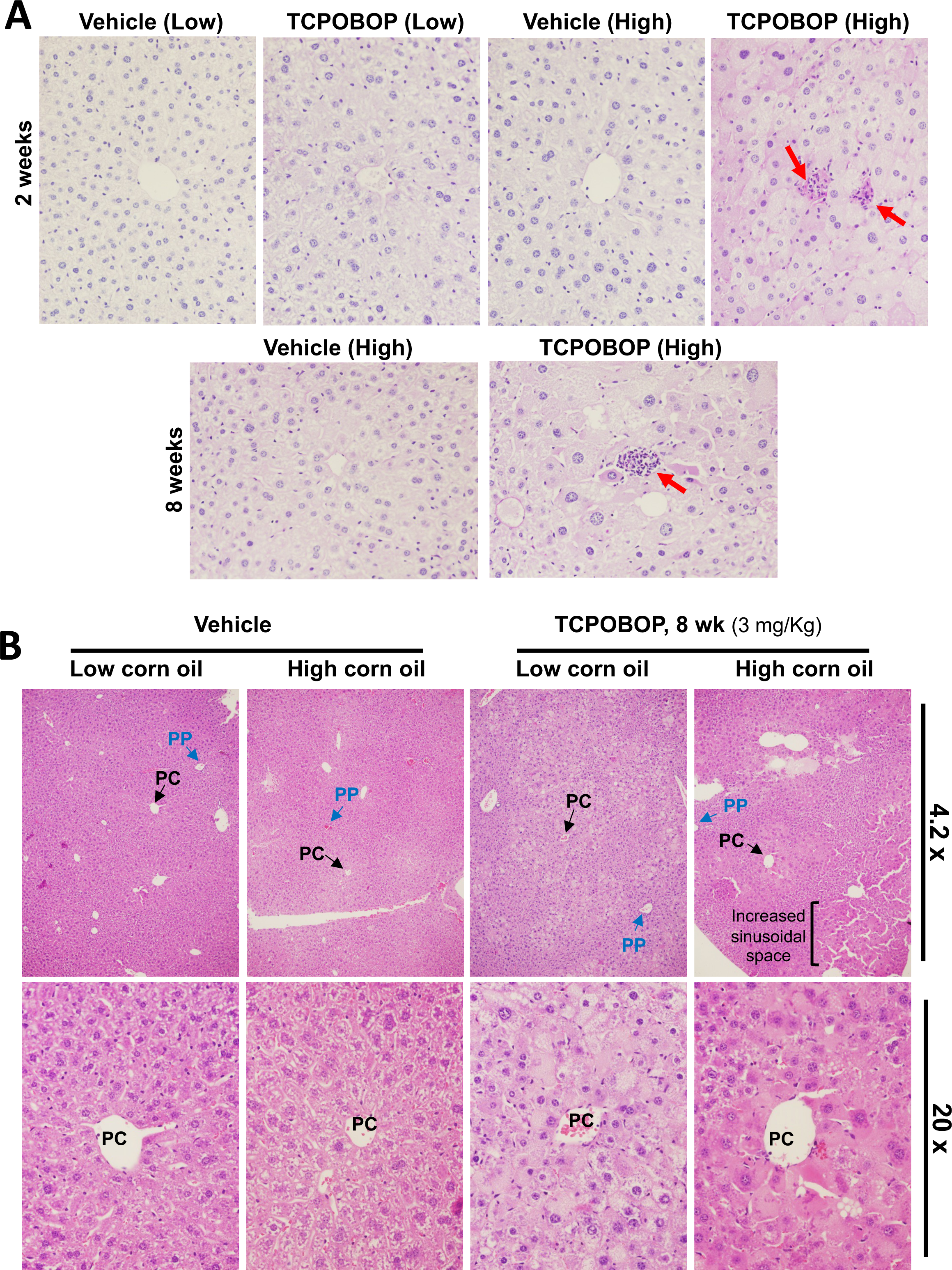
Increased focal immune cell infiltration and increased sinusoidal space induced by TCPOBOP/high corn oil regimen. **(A)** Diastase-stained liver sections reveal immune cell infiltration in livers from mice given TCPOBOP using the high corn oil vehicle regimen (‘High’) but not the low corn oil vehicle regimen (‘Low’) for 2 wk or for 8 wk, as indicated. The high corn oil vehicle regimen alone was without effect. **(B)** H&E staining reveals increase in sinusoidal space specifically in livers from mice given TCPOBOP using the high corn oil regimen. Also see Fig. S7.

RNA-seq analysis comparing TCPOBOP responses after 8 wk using the high vs. low corn oil regimen identified 870 genes showing higher expression with TCPOBOP when delivered with high corn oil, and 922 genes showing lower expression (Table S4A; FDR < 0.05). Top GO terms, related to ribosomes and mitochondria/mitochondrial respiratory chain, were highly enriched in the 8 wk TCPOBOP/high corn oil up regulated gene set (DAVID analysis; FDR = 4.45E-38, 6.85E-33, respectively), while metal ion binding, regulation of cell migration and regulation of RNA polymerase II transcription showed very strong enrichments in the 8 wk TCPOBOP/high corn oil down regulated gene set (FDR = 5.5E-10 to 3.2E-13) (Table S4C, Table S4D). Moreover, the top 3 IPA canonical pathways specifically activated by the 8 wk TCPOBOP high corn oil regimen, and not by the 2 wk TCPOBOP exposures, were EIF2 signaling, a critical stress-induced regulator of translation [116], oxidative phosphorylation and mitochondrial dysfunction (Fig. 4F), all key features of MASLD development and progression [117–119]. Other canonical pathways specific to the 8 wk TCPOBOP/high corn oil treatment group included mTOR signaling, a key regulator of lipid metabolism and of autophagy, which when dysregulated leads to liver diseases [120], as well as glucocorticoid and estrogen receptor signaling, phagosome maturation and BAG2 signaling (Fig. 4F).

To help elucidate mechanisms underlying the more advanced pathology induced by 8 wk TCPOBOP/high corn oil exposure, we identified upstream regulators specifically associated with this treatment (Fig. 5D). The top activated upstream regulator, MLXIPL, also known as carbohydrate-responsive element binding protein (ChREBP), is a carbohydrate-responsive transcription factor that activates genes of *de novo* lipogenesis and plays a key role in MASLD [121, 122]. Other activated upstream regulators specific to TCPOBOP/high corn oil treatment include Cab39l, a tumor suppressor that increases expression of mitochondrial respiration genes [123], and Ddx3x, a regulator of pro-survival stress granule assembly that protects hepatocytes from drug-induced liver injury [124]. TCPOBOP/high corn oil exposure inhibited the activities of four upstream regulators. One inhibited upstream regulator, Rictor, is a component of the mTOR signaling complex mTORC2, which has many functions, including controlling the balance of lipid and glucose in the liver [120, 125], while another inhibited upstream regulator, Larp1, is an RNA-binding protein that mediates specific translational regulation by the other major mTOR signaling complex, mTORC1 [126]. Finally, TCPOBOP/high corn oil treatment inhibited the function of the upstream regulator Kdm5a, a histone-H3 lysine-4 demethylase with oncogenic activity, whose knockdown suppresses liver cancer growth [127].

## Discussion

The NR transcription factor CAR can be activated by structurally diverse xenobiotics, including TCPOBOP, which dysregulates the expression of hundreds of liver-expressed genes within 3 h [40] and within a few days induces pronounced liver histopathological responses, including hypertrophy and hyperplasia [128]. Little is known, however, about the secondary gene responses induced by CAR activation and their associated hepatic pathological changes, which are expected to play an important role in the progression to hepatocellular tumors that emerge with high frequency in mice exposed to TCPOBOP persistently for 20-30 weeks [45, 46]. Here we address this gap by characterizing the pericentral liver steatosis and liver damage, and associated transcriptomic changes, that emerge by 2 wk after a single injection of TCPOBOP and, with continued TCPOBOP exposure, persist for at least 8 wk. Early (1 day) TCPOBOP-induced gene responses were enriched for genes of lipid and xenobiotic metabolism and protection from oxidative stress, and late (2 wk) responding genes, pathways and their TCPOBOP-activated upstream regulators expanded to encompass immune response/inflammatory disease, macrophage activation, and cytokine and reactive oxygen species production. Notable sex differences were observed in both the pathological and the transcriptomic changes that TCPOBOP induces, with steatosis in females being weaker in the pericentral region but stronger in the periportal region as compared to males, and with 2 wk TCPOBOP-activated upstream regulators in male but not female liver specifically enriched for terms such as defense response, cellular response to cytokine stimulus, DNA metabolic process and chromatin binding. Finally, more advanced liver pathology and unique upstream regulators associated with MASLD development and progression were induced when TCPOBOP was delivered using a high corn oil vehicle-based weekly injection regimen.

We used RNA-seq to develop a comprehensive picture of the time-dependent transcriptomic changes that TCPOBOP induces in mouse liver. Xenobiotic metabolism, as well as NRF2 oxidative stress response and glutathione-mediated detoxification, were significantly activated by 1 day, with upstream regulators involved in protection against oxidative stress, such as NFE2L2 (NRF2) and NFKBIA, being highly enriched in both sexes. However, other anti-oxidant upstream regulators, notably glutathione reductase (GSR) and thioredoxin reductase 1 (TXNRD1), were inhibited by TCPOBOP exposure, indicating an element of decreased protection from MASLD-associated oxidative stress. Many more genes were dysregulated after 2 wk TCPOBOP exposure than after 1 d in both sexes, due to expansion of the TCPOBOP response to include many indirect, secondary response genes, as evidenced by the absence of CAR binding at open chromatin regulatory sites at many of these genes. Consistent with this finding, the 2 wk/secondary TCPOBOP response genes included immune-related genes that are deficient in hepatocytes but preferentially expressed in liver non-parenchymal cells, where CAR expression is very low [41]. These late (2 wk) response genes and their immune-related upstream regulators (Fig. 5) are presumably activated as a secondary response to the hepatocyte damage that TCPOBOP induces as a primary response in hepatocytes. Examples of such liver non-parenchymal secondary response genes include *Gpnmb* and *Mmp12*, markers for MASH-associated macrophages induced in livers of high fat diet-fed mice [11, 80]; and *Col1a1*, a profibrogenic marker for activated, collagen-producing hepatic stellate cells [81]. Upstream regulators specifically activated at the 2 wk TCPOBOP time point included: STAT1, which promotes MASH [88, 89]; MYD88, which induces liver inflammation and M2 macrophage polarization [90, 91]; and several NFKB inflammatory pathway regulators [93]. Late upstream regulators, including the MASLD protective factors DUSP1 [94] and SIRT1 [95, 96], were predicted to be inhibited by TCPOBOP exposure, and may thus contribute to steatotic liver disease progression.

Macrophage activation and reactive oxygen species production and metabolism, and to a lesser extent inflammation, were identified as Disease and Bio Functions strongly enriched after 2 wk but not after 1 d TCPOBOP exposure. In contrast, the Disease and Bio Function hepatic steatosis was already enriched after 1 day, consistent with the early onset of steatosis seen by histological analysis. While lipid handling can be a protective, anti-lipotoxic function of macrophages in some contexts [129], hepatic macrophages integrate signals from steatotic hepatocytes together with systemic inflammation in a way that contributes to the progression from MASLD to MASH and fibrosis [130, 131]. We did not, however, observe hepatic fibrosis either 2 wk or 4 wk after initiating TCPOBOP treatment, indicating that the exposed livers have not yet transitioned from steatotic liver disease to a MASH-like stage, which presumably occurs later on during the progression of TCPOBOP-induced liver disease, which ultimately results in extensive liver tumors, as seen after 20-30 wk persistent TCPOBOP exposure [45, 46]. Nevertheless, liver fibrosis and liver damage were found to be top enriched IPA Tox Functions in both sexes at the 2 wk TCPOBOP time point (Table S5B); thus, gene signatures of fibrosis are already detectable within 2 wk of a single TCPOBOP injection.

CAR has complex effects and seemingly contradictory roles in steatotic liver disease. Activation of CAR induced pericentral steatosis within a few days of TCPOBOP injection in mice fed normal chow diet, as shown here, with prolonged TCPOBOP exposure activating gene programs and histopathologies associated with MASLD development, as discussed above. CAR-induced hepatic triglyceride accumulation in male mouse liver was previously associated with LXR-independent activation of several lipogenesis-related genes (*Elovl6*, *Fasn*, *Pklr*, *Thrsp*, *Pnpla3*, *Gck*) [132], all of which, except *Gck*, were confirmed here to be TCPOBOP induced (Fig. 3, Table S1B). Importantly, four of these genes (*Elovl6*, *Fasn*, *Pklr*, *Thrsp*) are key MASLD driver genes, as determined by integrative analysis of SNPs, gene expression and hepatic triglyceride datasets across the Hybrid Mouse Diversity panel [8]. CAR expression has also been closely linked with MALSD in human liver, where nuclear levels of CAR protein are significantly elevated in patients with steatohepatitis and are positively correlated with lipid droplet size [133]. Other studies demonstrate, however, that TCPOBOP-activated CAR can attenuate high fat diet-induced obesity and diabetes and improve hepatic steatosis [134, 135]. These steatotic liver disease protective effects of CAR are specifically manifested in the context of MASH-inducing high fat diets and are dependent on GADD45b [25], a CAR transcriptional co-activator that is itself highly induced in liver by TCPOBOP treatment [136]. Proposed mechanisms for the protection by CAR from diet-induced steatotic liver disease include repression of gluconeogenic gene expression by competition with HNF4 (Nr2a1) for binding to gene regulatory region sequences [134, 137, 138], suppression of PPARA-dependent fatty acid oxidation [139], and CAR-facilitated post-transcriptional ubiquitination leading to degradation of the transcriptional coactivator PGC-1α [140].

Sex differences in the incidence of MASLD (males > females) are associated with sex differences in the metabolism of lipids [60, 141, 142], drugs and steroids [143]. In particular, MASLD-associated steatosis and steatohepatitis are more severe in males, which have elevated levels of proinflammatory/profibrotic cytokines, and ultimately form liver tumors at a higher frequency than in females, as seen in both mouse models and humans [5–7]. CAR dysregulates liver gene expression in a sex-dependent manner (Fig. 4) [40], and late (2 wk) upstream regulators activated in male but not female liver were linked to pro-inflammatory responses and hepatocellular carcinoma progression. We also observed sex differences in the pattern of TCPOBOP-induced steatosis, with the pericentral pattern of TCPOBOP-induced neutral lipid accumulation stronger in male than in female liver, both at a saturating TCPOBOP dose (3 mg/kg) and at doses below saturation (0.2, 0.6 mg/kg) with respect to CAR activation [53]. In contrast, TCPOBOP stimulated greater periportal lipid accumulation in female than in male liver. This sex-dependent zonation of TCPOBOP-induced hepato-steatosis reflects the pericentral pattern of CAR expression across the liver lobule seen in male liver, when taken together with the higher basal level of CAR expression that females display in the periportal region, as revealed by single nucleus RNA-seq [41]. Importantly, these sex differences in CAR zonation and CAR-induced steatosis may have implications for the severity of liver pathology and therapeutic outcomes, which differ between periportal and pericentral liver disease [144]. Together, these findings support the proposal that the level of CAR expression is a limiting factor in the pathological response to TCPOBOP and its zonation, consistent with the positive correlation reported between CAR levels and steatohepatitis across a panel of human livers [133].

Finally, we observed more extensive liver pathology and associated transcriptomic changes when weekly TCPOBOP injections were given using a high corn oil vehicle. This TCPOBOP exposure regimen activated several pathways and responses key to MASLD development and progression [117–119], including mitochondrial dysfunction, a mechanistic driver of MASLD [8], and EIF2 signaling, a critical stress-induced regulator of translation [116]. The mechanistic basis for the increased pathology seen when TCPOBOP was given via a high corn oil vehicle is unknown, but likely involves one or more of the unique upstream regulators that we identified for this treatment regimen. One such activated upstream regulator is the carbohydrate-responsive transcription factor MLXIPL, which induces genes of *de novo* lipogenesis and plays a key role in MASLD [121, 122]; and one of the inhibited upstream regulators, RICTOR, is a central component of the mTOR signaling complex mTORC2, which regulates the balance between lipid and glucose in the liver [120, 125].

In conclusion, the 2 wk TCPOBOP exposure (single injection) mouse liver model recapitulates several early phenotypes of high fat diet-induced MASLD, including pericentral steatosis and hepatocyte damage. It also induces widespread, secondary gene expression changes that apparently involve liver non-parenchymal cells, consistent with the emergence of other key features of MASLD. TCPOBOP also induces what appear to be early gene signatures indicative of the transition to a MASH-like state, including inflammatory response, macrophage activation and liver fibrosis genes. TCPOBOP-treated mice may thus serve as a useful model for further investigation of the mechanisms by which foreign chemicals induce foreign chemical-dependent MASLD development and the subsequent transition from MASLD to MASH. Upstream regulators that were either activated or repressed by TCPOBOP were identified, some of which could be useful markers for the transition from MASLD to MASH and may potentially serve as targets for reversing disease progression.

## Abbreviations

Ahr: aryl hydrocarbon receptor
CAR: constitutive androstane receptor, *Nr1i3*
HCC: hepatocellular carcinoma
IPA: Ingenuity Pathway Analysis
MASH: metabolic dysfunction-associated steatohepatitis
MASLD: metabolic dysfunction-associated steatotic liver disease
NR: nuclear receptor
PXR: pregnane X receptor
TCPOBOP: (1,4-bis[2-(3,5-dichloropyridyloxy)]benzene).

## Conflicts of interest

The authors declare that they have no conflicts of interest.

## Funding

Supported in part by NIH grant ES024421 (to DJW).

## Author contributions

All animal experiments and other wet lab analyses were carried out by HM and RS. Histology analysis and preparation of related figures and associated statistical analyses were primarily performed by RS. qPCR analysis, RNA-seq library preparation and initial data analysis and related figure preparation were performed by HM. All other data analysis was performed by DJW. DJW provided guidance and supervised the overall project. The manuscript was drafted by DJW with input from HM and the final manuscript was edited by DJW. All authors reviewed and approved of the final manuscript.

## Supplementary Figure legends

**Fig. S1.**
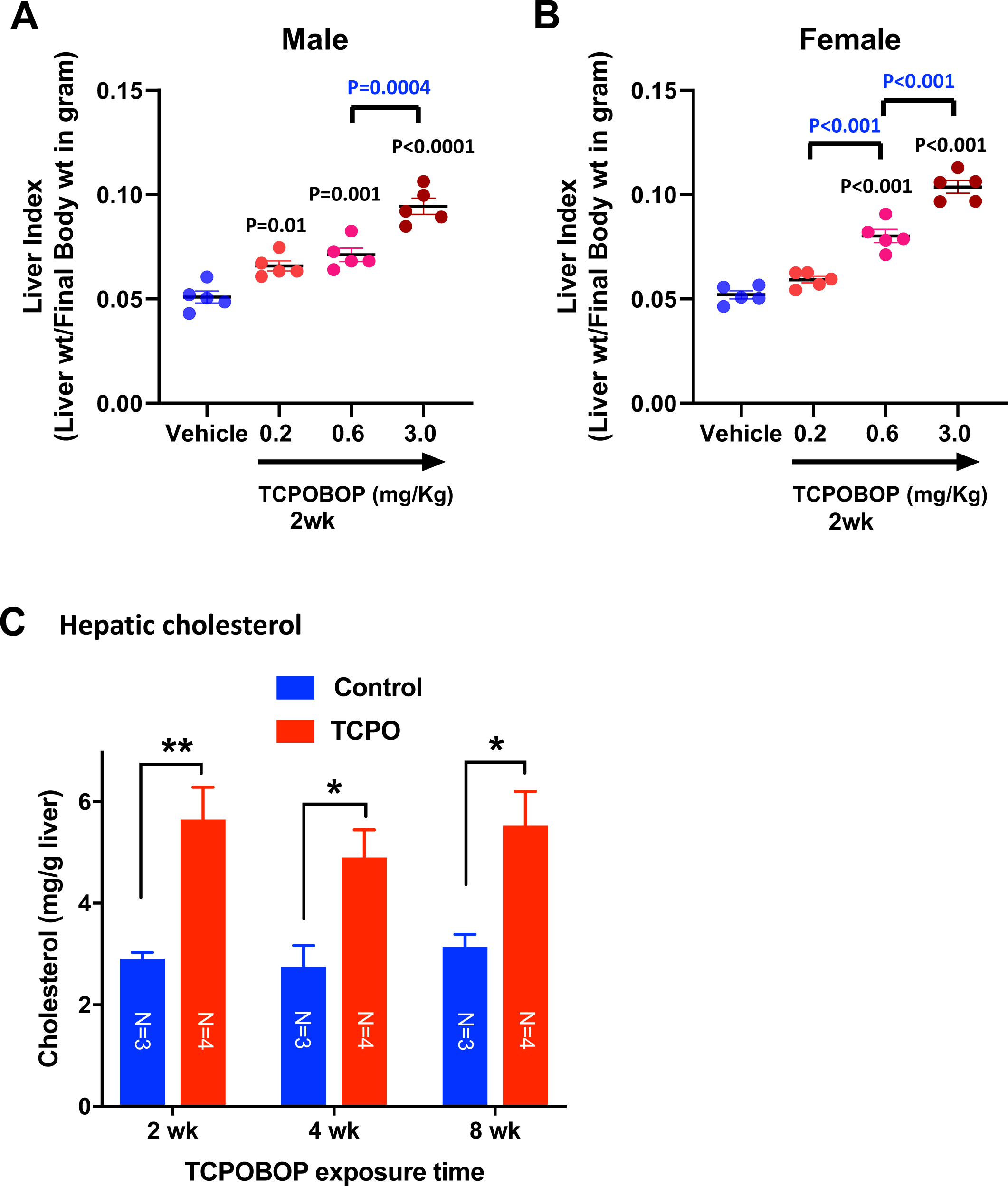
**(A, B)** Liver index for dose-response of TCPOBOP exposure in male and female mouse liver. Mice were given a single injection of TCPOBOP at the doses indicated (low corn oil regimen); livers were collected 2 wk later. Data shown are mean +/- SEM for n=5 livers/group. Significance values in *black* are for comparisons to vehicle controls; those in *blue* are for comparisons between adjacent doses, as indicated. **(C)** TCPOBOP increases hepatic cholesterol content in livers of male mice (n=3-4/livers group) given weekly injections of TCPOBOP using the high corn oil regimen (see Methods), with livers collected after 2, 4 or 8 wk. Two-way ANOVA with Tukey’s multiple comparisons test comparing TCPOBOP exposure to male or female vehicle controls.

**Fig. S2.**
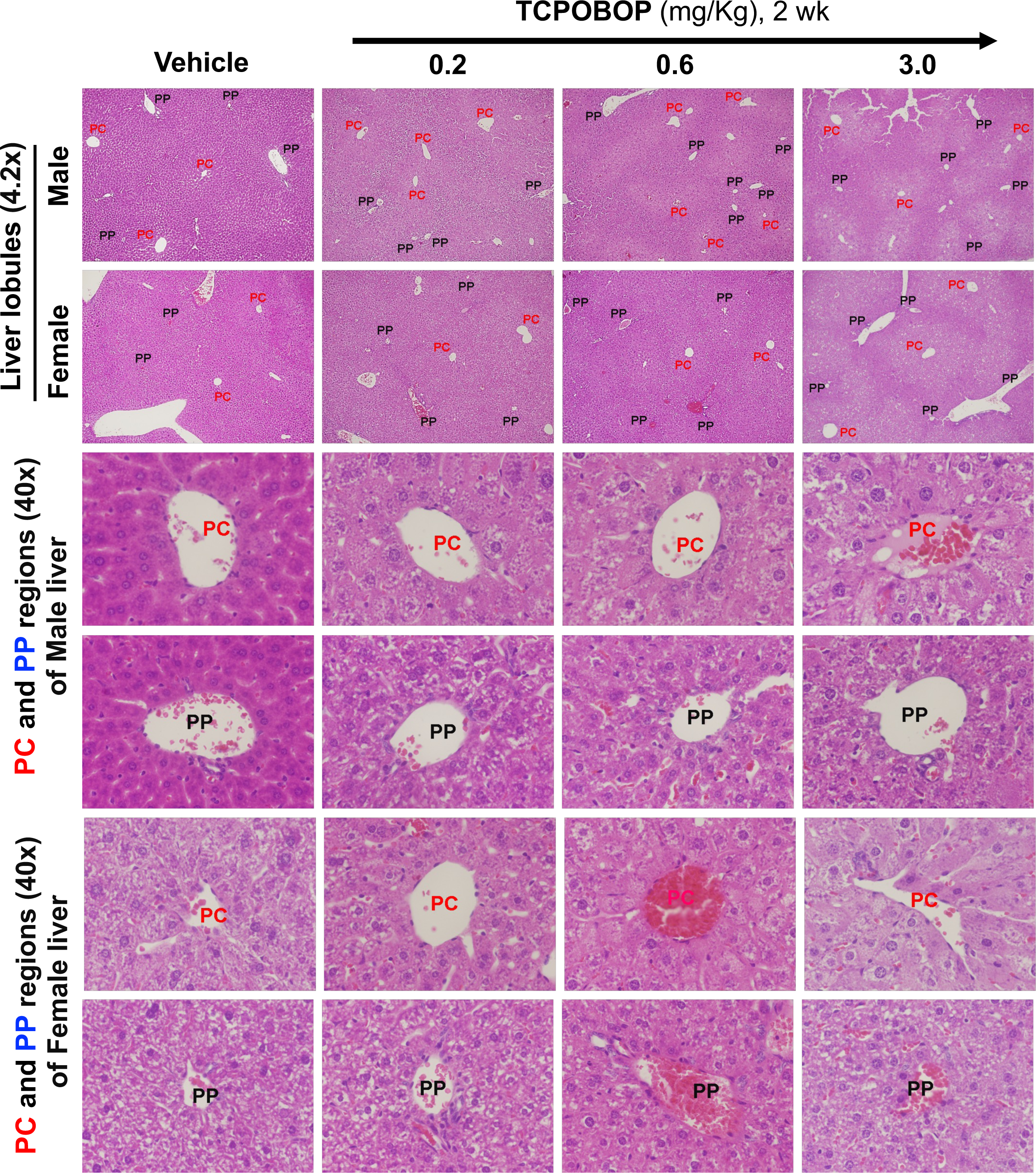
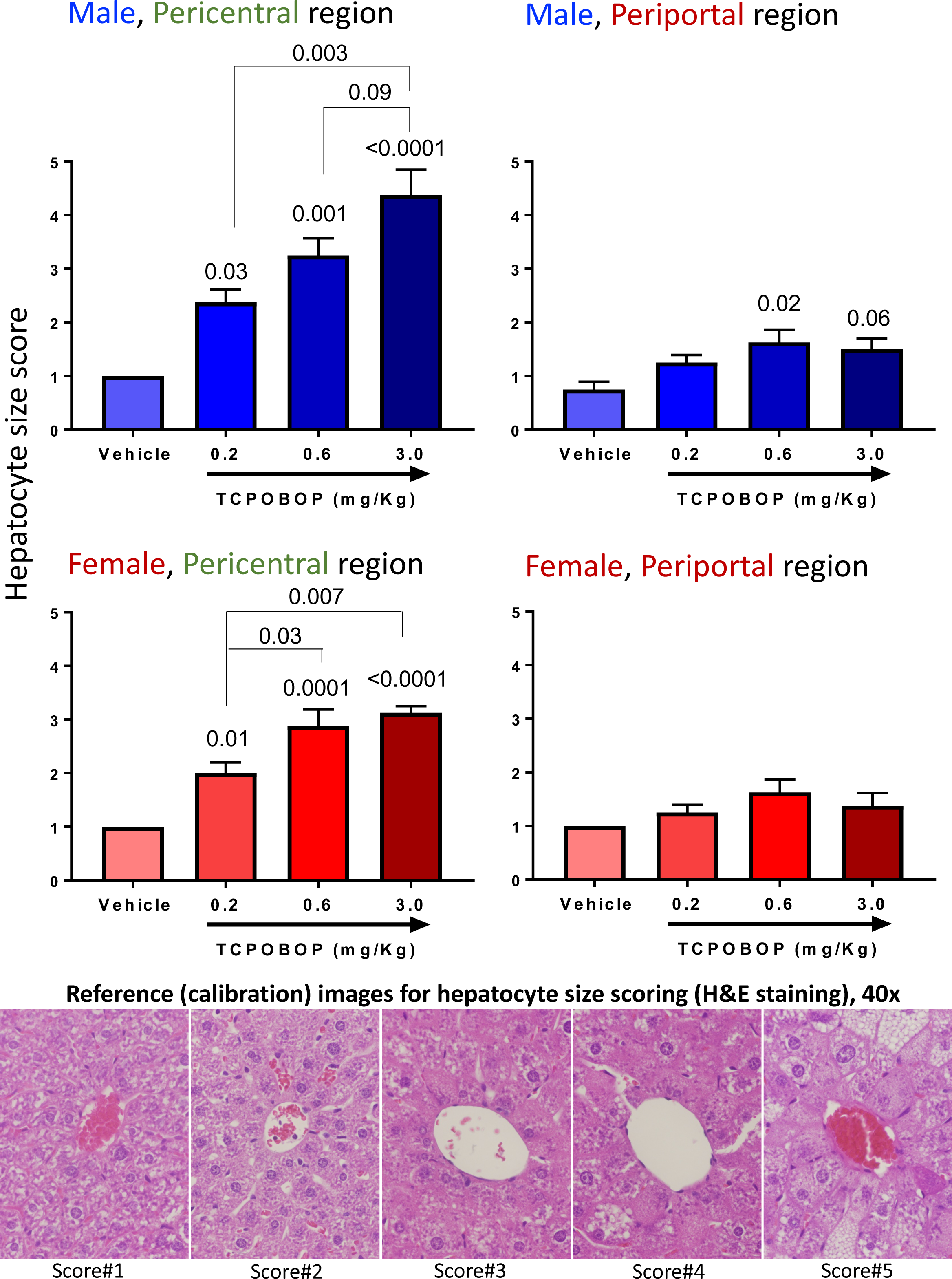
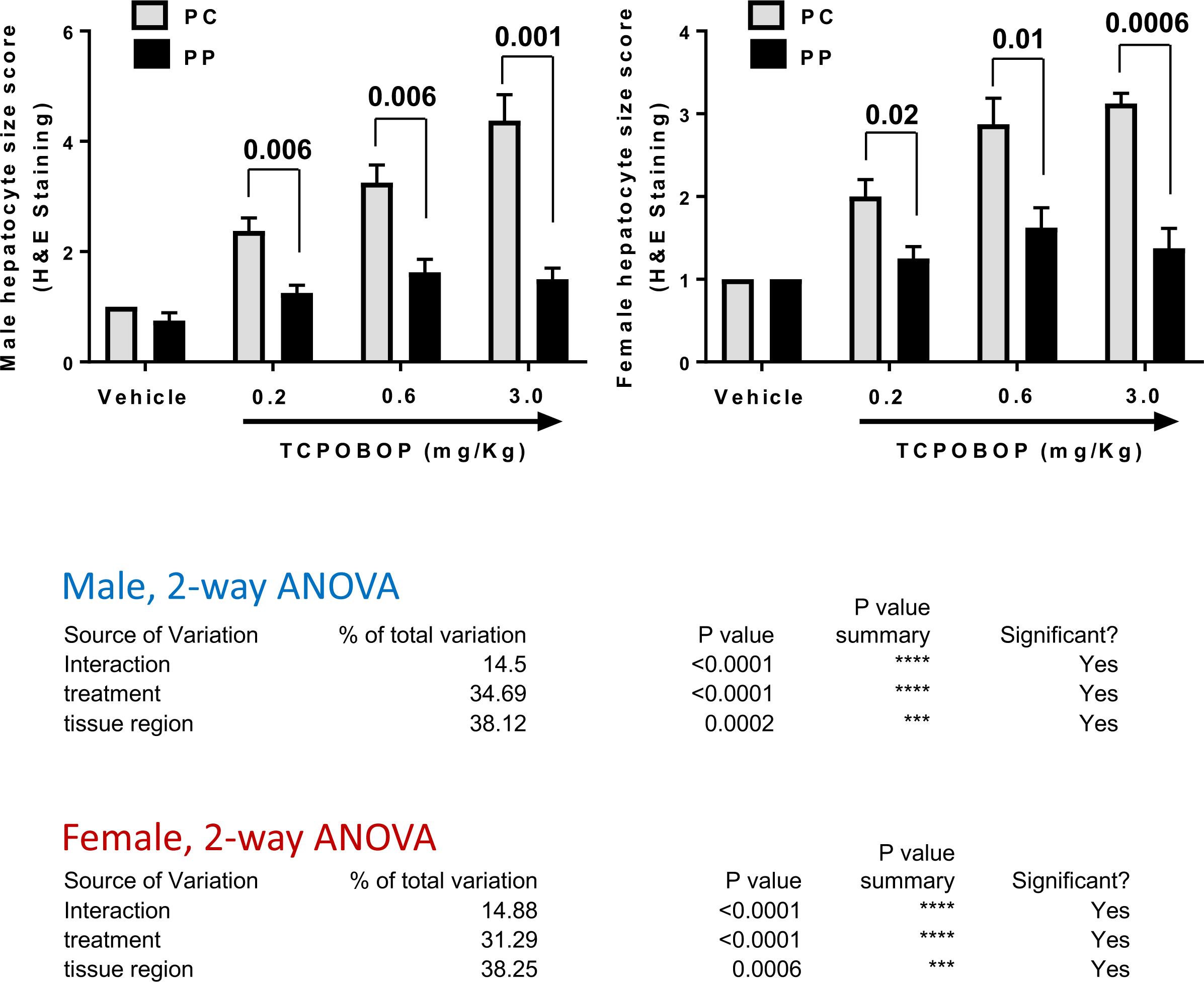
**(A)** H&E staining of livers from male and female mice given a single injection of TCPOBOP or vehicle control at the doses indicated (low corn oil regimen); livers were collected 2 wk later. Representative images (n=5 livers/group) are shown for both the periportal (PP) and pericentral (PC) liver lobule regions, as marked. Livers are from the same group of mice shown in Fig. 2 and in Fig. S1A-S1B, with imaging at either 4.2x (top) or 40x, as marked. **(B)** Hepatocyte size scoring for periportal and pericentral male and female hepatocytes from 2 wk TCPOBOP dose-response study shown in (A). Reference images used for scoring are shown at the bottom. Data shown are mean +/- SEM for n=5 livers/group, with statistical significance compared to the vehicle control determined by Tukey’s multiple comparisons test (values directly above each bar). **(C)** Comparison of hepatocyte size between pericentral and periportal regions in male livers (*left*) and in female livers (*right*). Data shown are mean +/- SEM for n=5 livers/group, with statistical significance compared to the vehicle control determined by t-test (values directly above each bar). Significance was also assessed by 2-way ANOVA (bottom).

**Fig. S3.**
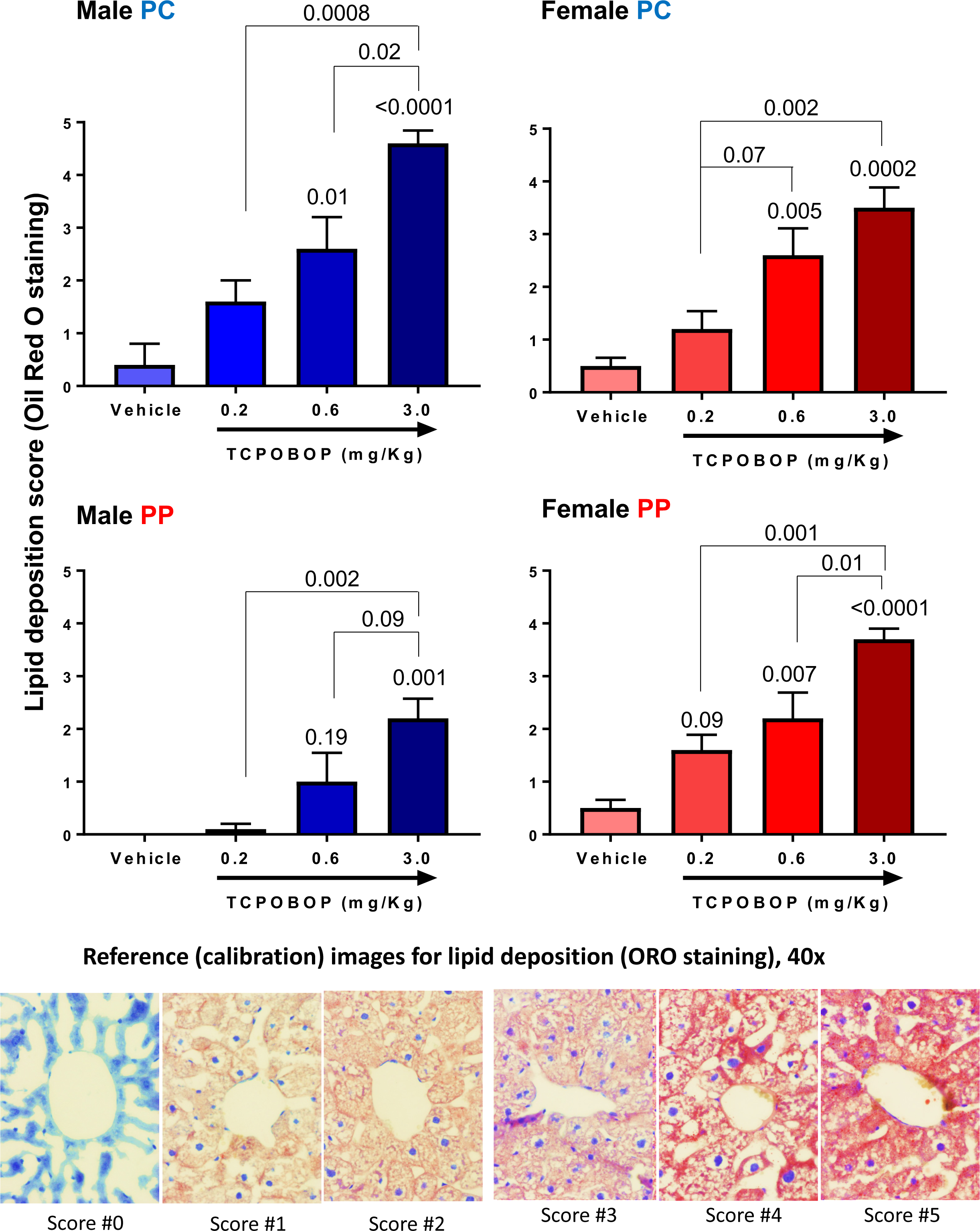
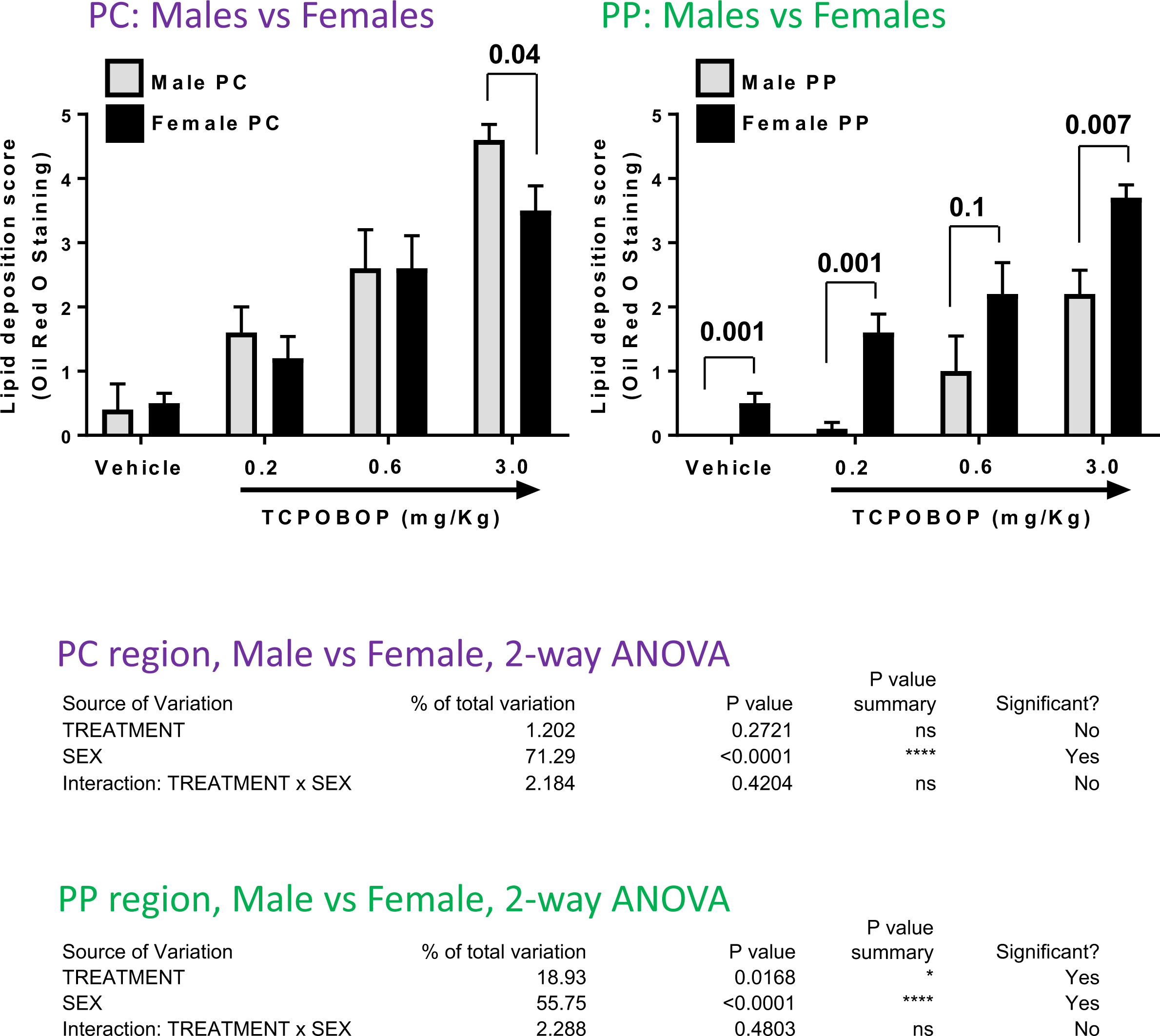
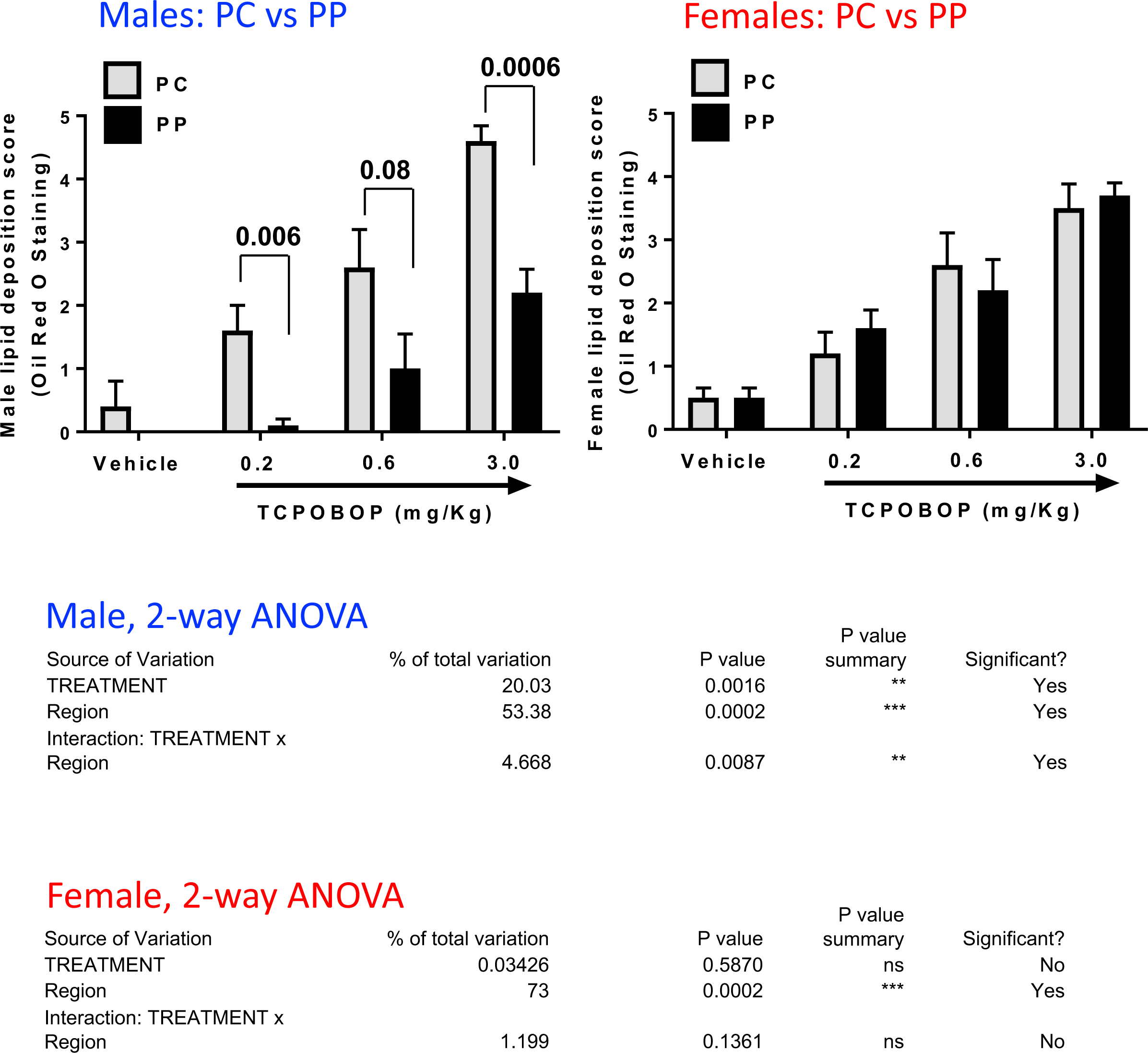
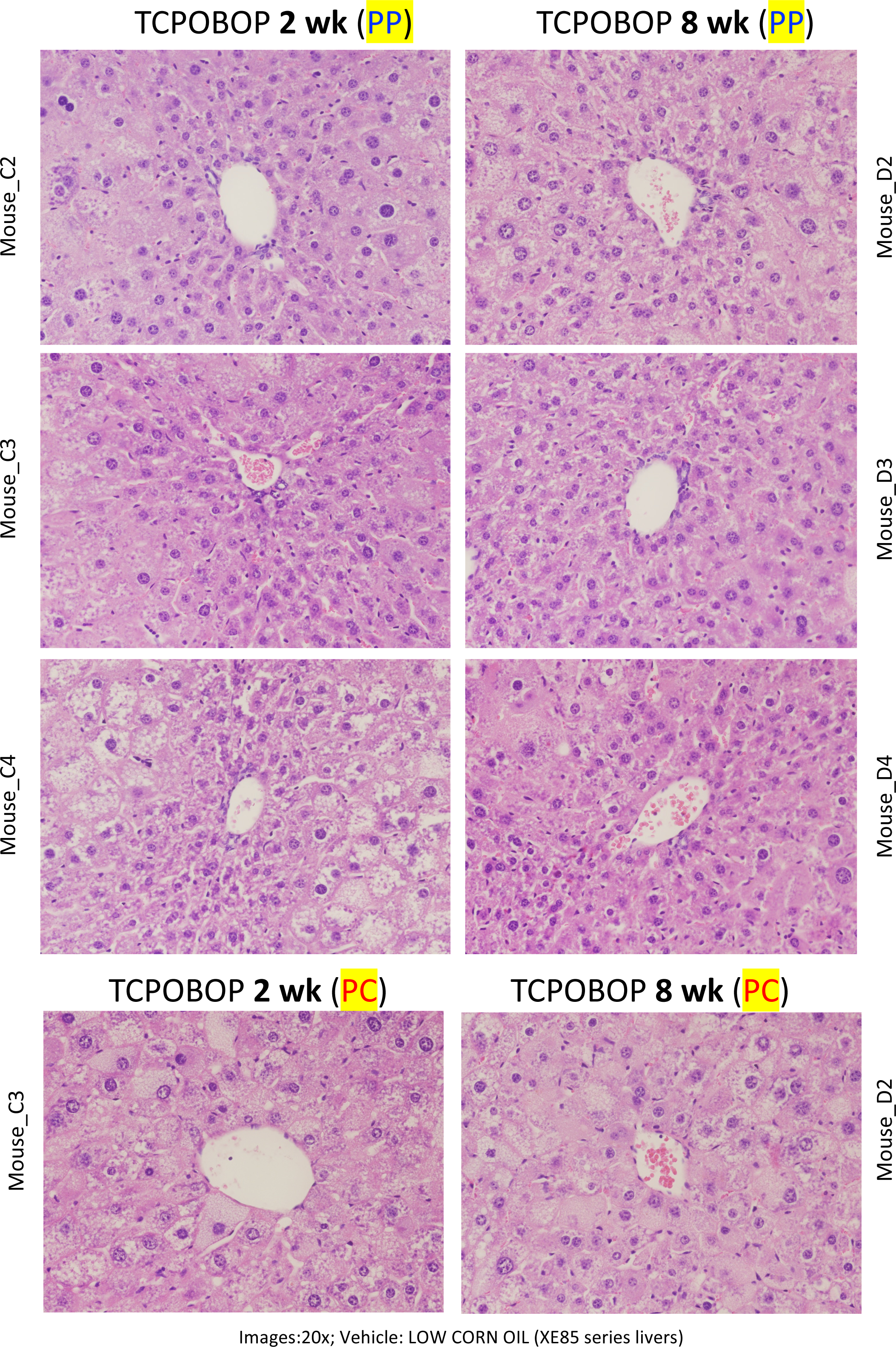
**(A)** Oil Red O staining intensity for pericentral (PC) and periportal (PP) male and female hepatocytes from the 2 wk TCPOBOP dose-response livers shown in Fig. 2. Reference images used for scoring are shown at the bottom. Data shown are mean +/- SEM for n=5 livers/group, with statistical significance compared to the vehicle control determined by Tukey’s multiple comparisons test (values directly above each bar). Also shown are those comparisons between TCPOBOP doses that are statistically different from each other, as marked by horizontal brackets above adjacent bars. **(B)** Comparison of Oil Red O staining intensity between males and females in the pericentral region (*left*) and in the periportal region (*right*). Data shown are mean +/- SEM for n=5 livers/group, with statistical significance compared to the vehicle control determined by t-test (values directly *above* each bar). Significance was also assessed by 2-way ANOVA (*bottom*). **(C)** Comparison of Oil Red O staining intensity between pericentral and periportal regions. Data shown are mean +/- SEM for n=5 livers/group, with statistical significance compared to the vehicle control determined by t-test (values directly *above* each bar). Significance was also assessed by 2-way ANOVA (*bottom*). **(D)** H&E stained sections reveal a similar degree of male mouse liver pathology after 2 wk vs after 8 wk TCPOBOP treatment (low corn oil regimen). Images at top show the periportal region, at the center of each image, as well as more distant, mid-lobular hepatocytes, which generally show greater pathology (increased hepatocyte size, increased lipid accumulation) as compared to the more immediate layers of periportal cells. Pericentral region is shown at the bottom for one liver from each group. Shown are representative images from n=4 livers per time point.

**Fig. S4.**
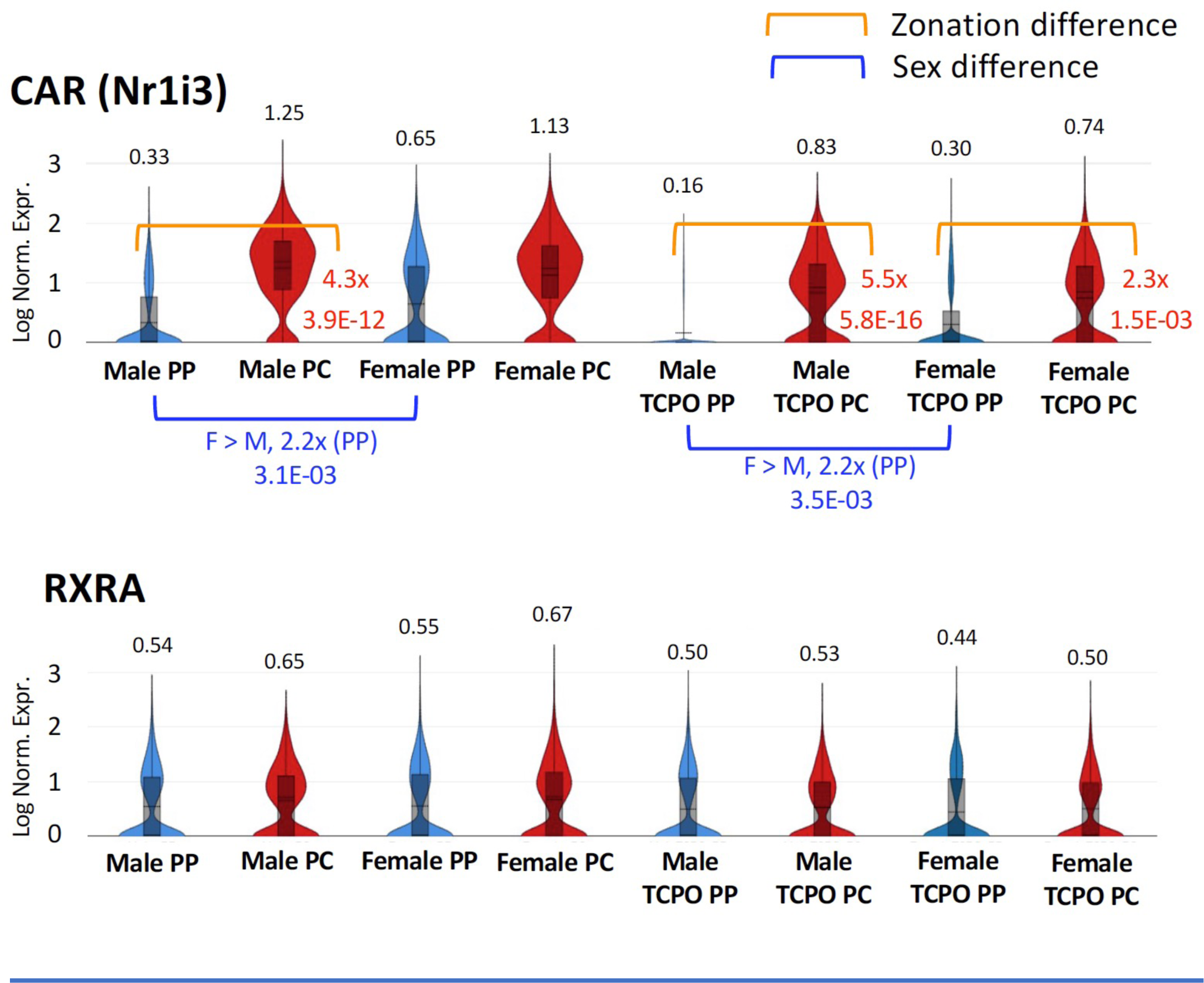
Single nucleus RNA-seq analysis showing zonation of CAR (Nr1i3) and RXRA expression in male and female mouse liver. Data are presented as violin plots, with mean values shown above each plot. Horizontal brackets mark significant differences in expression and their associated fold-change values between cell populations. Orange bracket, significant zonation differences; Blue bracket, significant sex differences. TCPO, 1 d exposure to TCPOBOP at 3 mg/kg, low corn oil regimen. Absence of bracket indicates no significant difference in zonation or sex bias. Data shown are from Goldfarb et al (2022), PMID: 35512247; DOI: 10.1210/endocr/bqac059.

**Fig. S5.**
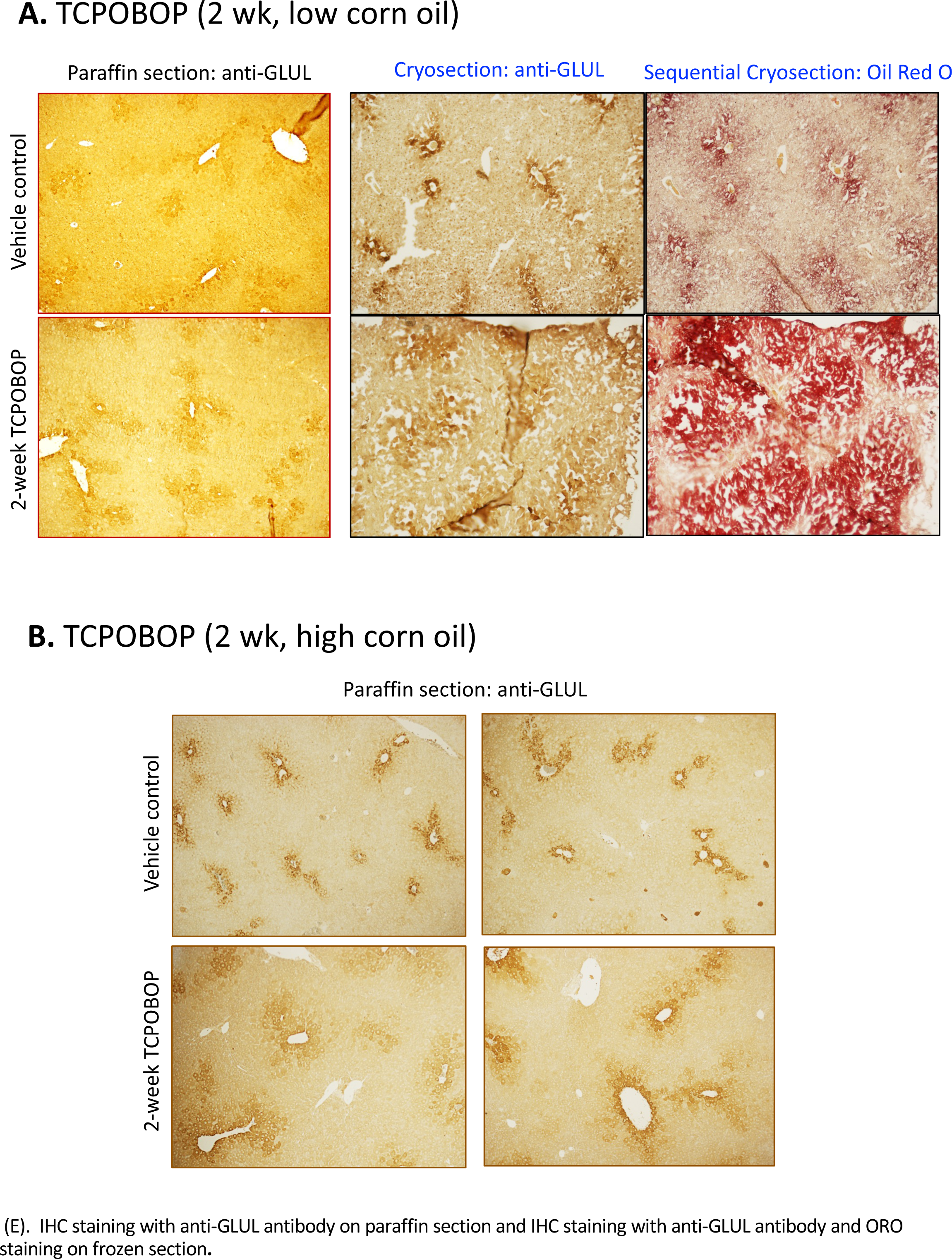
Immunostaining with anti-GLUL (glutamate-ammonia ligase, i.e., glutamine synthase, an established marked for pericentral hepatocytes) applied to paraffin sections or cryosections (as indicated) from male mouse livers. Sections were prepared from vehicle control or from 2 wk TCPOBOP-treated livers, delivered using the low corn oil regimen **(A)** or the high corn oil regimen **(B)**. GLUL immunostaining is highly localized around the central vein in vehicle control livers but is more diffuse and encompasses a few additional cell layers more distant from the central vein in TCPOBOP-exposed livers. The same TCPOBOP-induced shift in GLUL staining pattern is seen with both vehicle control regimens (A vs B). The cryosections in (A) (*blue* text) are sequential sections from the same liver that were stained with anti-GLUL antibody and with Oil Red O, respectively, to highlight the localization of Oil Red O staining to the GLUL-immunostained pericentral region. Anti-GLUL immunostained sections in (B) were obtained from 2 individual livers per vehicle control or TCPOBOP-treated group.

**Fig. S6.**
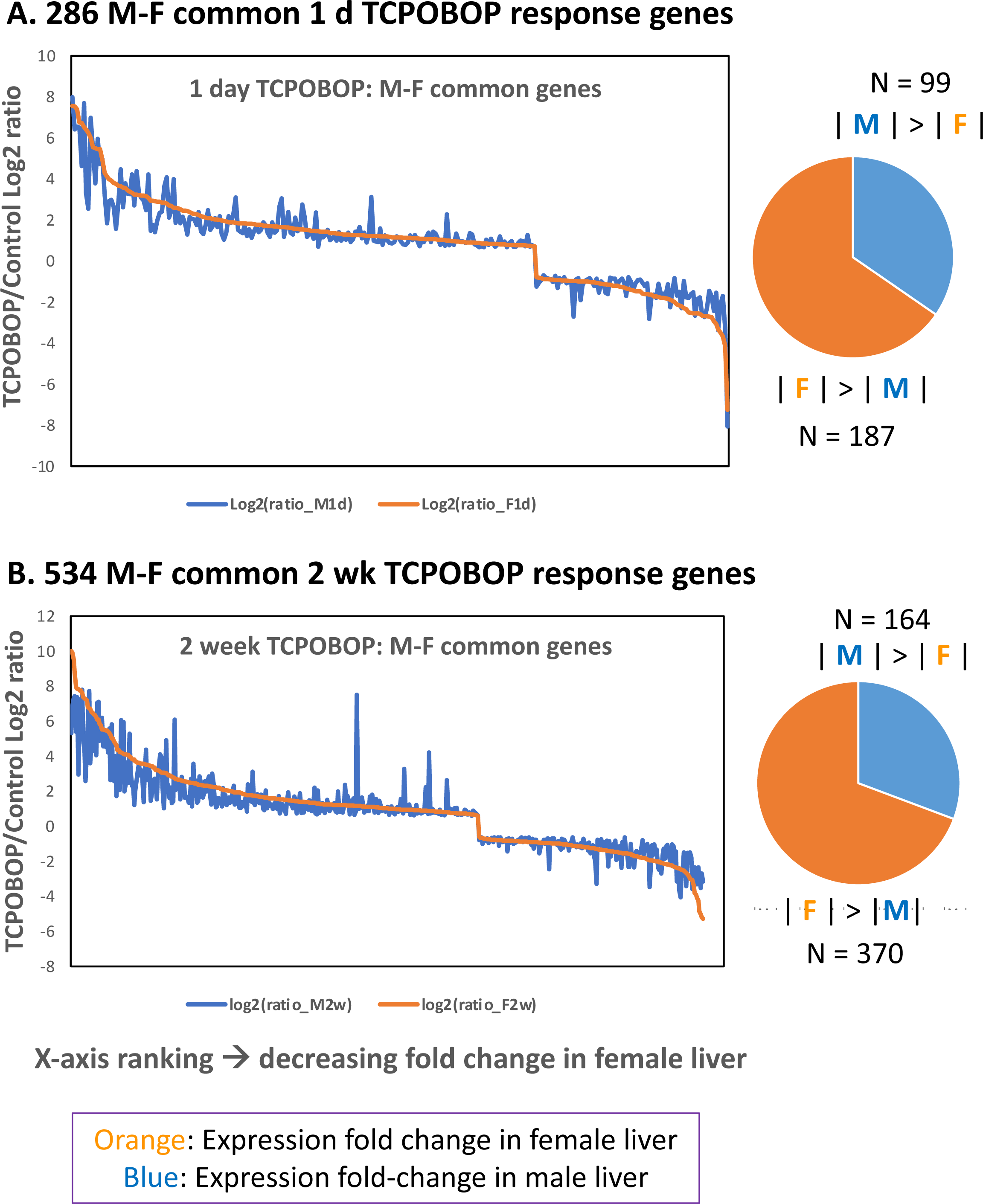
Genes that respond to TCPOBOP in both male and female mouse liver more often show greater fold-changes in expression in female liver. Graphs show log2 of gene expression ratios (TCPOBOP/Vehicle control) for genes that respond to TCPOBOP treatment in the same direction in male (*blue*) as in female (*orange*) livers at |fold-change| > 1.5 and FDR < 0.05. Genes are ranked along the X-axis by their decreasing log2 TCPOBOP/Vehicle ratios in female liver. Pie charts at the right show the proportion of genes at each time point, and the number of genes whose response to TCPOBOP (expression fold change) is greater in either male liver, or in female liver, as marked. **(A)** 286 genes that respond in common in both sexes to 1 day TCPOBOP treatment. **(B)** 534 genes that respond in common in both sexes to 2 wk TCPOBOP treatment. A small number of genes that responded to TCPOBOP in both sexes, but in opposite direction, were omitted.

**Fig. S7.**
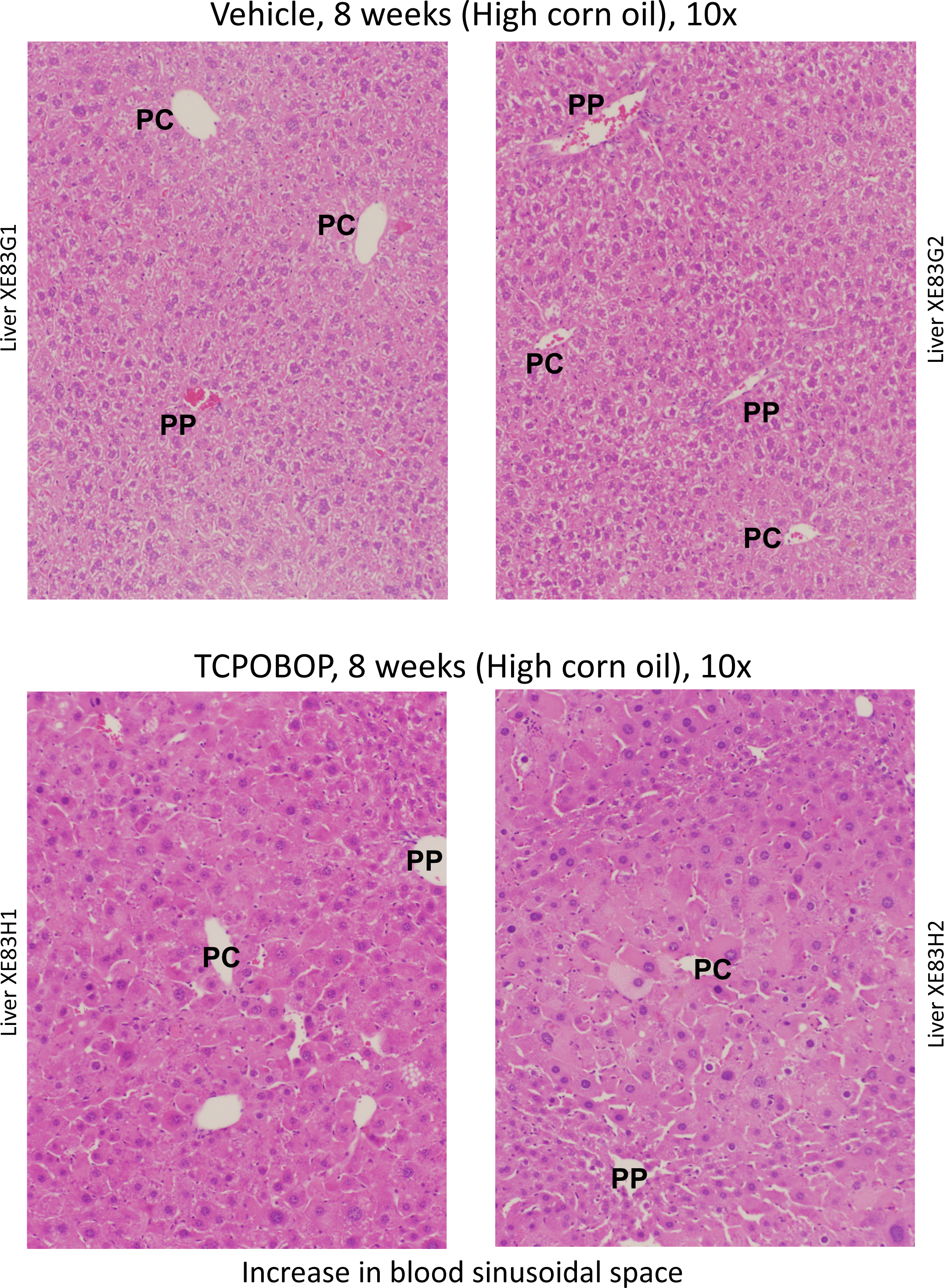
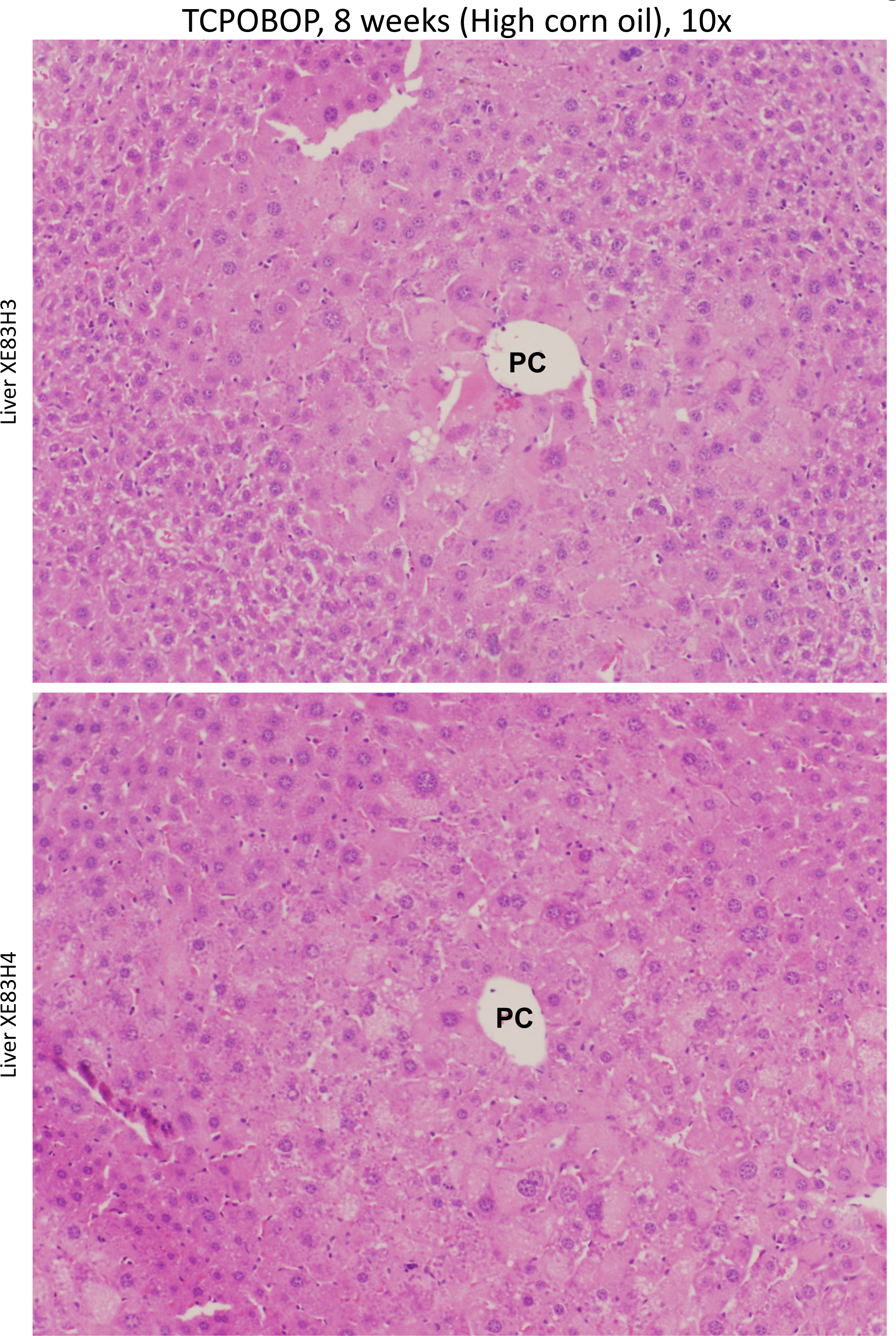
Liver sections stained with H&E from 4 individual mice from the 8 week TCPOBOP/high corn oil regimen treatment group. Regions with an increase in blood sinusoidal space are seen in each liver from the TCPOBOP-treated mice.

## Notes

### Competing Interest Statement

The authors have declared no competing interest.

## References

[1] M. Eslam, P.N. Newsome, S.K. Sarin, Q.M. Anstee, G. Targher, M. Romero-Gomez, S. Zelber-Sagi, V. Wai-Sun Wong, J.F. Dufour, J.M. Schattenberg, T. Kawaguchi, M. Arrese, L. Valenti, G. Shiha, C. Tiribelli, H. Yki-Järvinen, J.G. Fan, H. Grønbæk, Y. Yilmaz, H. Cortez-Pinto, C.P. Oliveira, P. Bedossa, L.A. Adams, M.H. Zheng, Y. Fouad, W.K. Chan, N. Mendez-Sanchez, S.H. Ahn, L. Castera, E. Bugianesi, V. Ratziu, J. George, A new definition for metabolic dysfunction-associated fatty liver disease: An international expert consensus statement, J Hepatol, 73 (2020) 202–209.

[2] Y. Takahashi, E. Dungubat, H. Kusano, T. Fukusato, Pathology and Pathogenesis of Metabolic Dysfunction-Associated Steatotic Liver Disease-Associated Hepatic Tumors, Biomedicines, 11 (2023).

[3] S. Wang, S.L. Friedman, Found in translation-Fibrosis in metabolic dysfunction-associated steatohepatitis (MASH), Sci Transl Med, 15 (2023) eadi0759.

[4] D. Kaltenecker, M. Themanns, K.M. Mueller, K. Spirk, T. Suske, O. Merkel, L. Kenner, A. Luís, A. Kozlov, J. Haybaeck, M. Müller, X. Han, R. Moriggl, Hepatic growth hormone - JAK2 - STAT5 signalling: Metabolic function, non-alcoholic fatty liver disease and hepatocellular carcinoma progression, Cytokine, 124 (2019) 154569.

[5] A. Lonardo, F. Nascimbeni, S. Ballestri, D. Fairweather, S. Win, T.A. Than, M.F. Abdelmalek, A. Suzuki, Sex Differences in Nonalcoholic Fatty Liver Disease: State of the Art and Identification of Research Gaps, Hepatology, 70 (2019) 1457–1469.

[6] X.Y. Chen, C. Wang, Y.Z. Huang, L.L. Zhang, Nonalcoholic fatty liver disease shows significant sex dimorphism, World J Clin Cases, 10 (2022) 1457–1472.

[7] R. Nevola, G. Tortorella, V. Rosato, L. Rinaldi, S. Imbriani, P. Perillo, D. Mastrocinque, M. La Montagna, A. Russo, G. Di Lorenzo, M. Alfano, M. Rocco, C. Ricozzi, K. Gjeloshi, F.C. Sasso, R. Marfella, A. Marrone, L.A. Kondili, N. Esposito, E. Claar, D. Cozzolino, Gender Differences in the Pathogenesis and Risk Factors of Hepatocellular Carcinoma, Biology (Basel), 12 (2023).

[8] K. Chella Krishnan, Z. Kurt, R. Barrere-Cain, S. Sabir, A. Das, R. Floyd, L. Vergnes, Y. Zhao, N. Che, S. Charugundla, H. Qi, Z. Zhou, Y. Meng, C. Pan, M.M. Seldin, F. Norheim, S. Hui, K. Reue, A.J. Lusis, X. Yang, Integration of Multi-omics Data from Mouse Diversity Panel Highlights Mitochondrial Dysfunction in Non-alcoholic Fatty Liver Disease, Cell Syst, 6 (2018) 103–115.e107.

[9] A. Loft, A.J. Alfaro, S.F. Schmidt, F.B. Pedersen, M.K. Terkelsen, M. Puglia, K.K. Chow, A. Feuchtinger, M. Troullinaki, A. Maida, G. Wolff, M. Sakurai, R. Berutti, B. Ekim Üstünel, P. Nawroth, K. Ravnskjaer, M.B. Diaz, B. Blagoev, S. Herzig, Liver-fibrosis-activated transcriptional networks govern hepatocyte reprogramming and intra-hepatic communication, Cell Metab, 33 (2021) 1685–1700.e1689.

[10] H. Yang, M. Arif, M. Yuan, X. Li, K. Shong, H. Türkez, J. Nielsen, M. Uhlén, J. Borén, C. Zhang, A. Mardinoglu, A network-based approach reveals the dysregulated transcriptional regulation in non-alcoholic fatty liver disease, iScience, 24 (2021) 103222.

[11] K. Karri, D.J. Waxman, Dysregulation of murine long noncoding single-cell transcriptome in nonalcoholic steatohepatitis and liver fibrosis, Rna, 29 (2023) 977–1006.

[12] A. Mosca, M. Manco, M.R. Braghini, S. Cianfarani, G. Maggiore, A. Alisi, A. Vania, Environment, Endocrine Disruptors, and Fatty Liver Disease Associated with Metabolic Dysfunction (MASLD), Metabolites, 14 (2024).

[13] A. Dolce, S. Della Torre, Sex, Nutrition, and NAFLD: Relevance of Environmental Pollution, Nutrients, 15 (2023).

[14] A. Aguayo-Orozco, F.Y. Bois, S. Brunak, O. Taboureau, Analysis of Time-Series Gene Expression Data to Explore Mechanisms of Chemical-Induced Hepatic Steatosis Toxicity, Front Genet, 9 (2018) 396.

[15] B. Wahlang, RISING STARS: Sex differences in toxicant-associated fatty liver disease, J Endocrinol, 258 (2023).

[16] B. Wahlang, J. Jin, J.I. Beier, J.E. Hardesty, E.F. Daly, R.D. Schnegelberger, K.C. Falkner, R.A. Prough, I.A. Kirpich, M.C. Cave, Mechanisms of Environmental Contributions to Fatty Liver Disease, Curr Environ Health Rep, 6 (2019) 80–94.

[17] M.C. Cave, H.B. Clair, J.E. Hardesty, K.C. Falkner, W. Feng, B.J. Clark, J. Sidey, H. Shi, B.A. Aqel, C.J. McClain, R.A. Prough, Nuclear receptors and nonalcoholic fatty liver disease, Biochim Biophys Acta, 1859 (2016) 1083–1099.

[18] J. Jin, B. Wahlang, H. Shi, J.E. Hardesty, K.C. Falkner, K.Z. Head, S. Srivastava, M.L. Merchant, S.N. Rai, M.C. Cave, R.A. Prough, Dioxin-like and non-dioxin-like PCBs differentially regulate the hepatic proteome and modify diet-induced nonalcoholic fatty liver disease severity, Med Chem Res, 29 (2020) 1247–1263.

[19] P. Honkakoski, T. Sueyoshi, M. Negishi, Drug-activated nuclear receptors CAR and PXR, Ann Med, 35 (2003) 172–182.

[20] D.J. Waxman, P450 gene induction by structurally diverse xenochemicals: central role of nuclear receptors CAR, PXR, and PPAR, Arch Biochem Biophys, 369 (1999) 11–23.

[21] B. Mackowiak, J. Hodge, S. Stern, H. Wang, The Roles of Xenobiotic Receptors: Beyond Chemical Disposition, Drug Metab Dispos, 46 (2018) 1361–1371.

[22] J. Wang, P. Lu, W. Xie, Atypical functions of xenobiotic receptors in lipid and glucose metabolism, Med Rev (Berl), 2 (2022) 611–624.

[23] L. Rakateli, R. Huchzermeier, E.P.C. van der Vorst, AhR, PXR and CAR: From Xenobiotic Receptors to Metabolic Sensors, Cells, 12 (2023).

[24] F. Oliviero, W. Klement, L. Mary, Y. Dauwe, Y. Lippi, C. Naylies, V. Gayrard, N. Marchi, L. Mselli-Lakhal, CAR Protects Females from Diet-Induced Steatosis and Associated Metabolic Disorders, Cells, 12 (2023).

[25] X. Cai, Y. Feng, M. Xu, C. Yu, W. Xie, Gadd45b is required in part for the anti-obesity effect of constitutive androstane receptor (CAR), Acta Pharm Sin B, 11 (2021) 434–441.

[26] M. Negishi, K. Kobayashi, T. Sakuma, T. Sueyoshi, Nuclear receptor phosphorylation in xenobiotic signal transduction, J Biol Chem, 295 (2020) 15210–15225.

[27] I. Tzameli, P. Pissios, E.G. Schuetz, D.D. Moore, The xenobiotic compound 1,4-bis[2-(3,5-dichloropyridyloxy)]benzene is an agonist ligand for the nuclear receptor CAR, Mol Cell Biol, 20 (2000) 2951–2958.

[28] S. Mutoh, M. Sobhany, R. Moore, L. Perera, L. Pedersen, T. Sueyoshi, M. Negishi, Phenobarbital indirectly activates the constitutive active androstane receptor (CAR) by inhibition of epidermal growth factor receptor signaling, Sci Signal, 6 (2013) ra31.

[29] A. Rampersaud, N.J. Lodato, A. Shin, D.J. Waxman, Widespread epigenetic changes to the enhancer landscape of mouse liver induced by a specific xenobiotic agonist ligand of the nuclear receptor CAR, Toxicol Sci, 171 (2019) 315–338.

[30] B. Niu, D.M. Coslo, A.R. Bataille, I. Albert, B.F. Pugh, C.J. Omiecinski, In vivo genome-wide binding interactions of mouse and human constitutive androstane receptors reveal novel gene targets, Nucleic Acids Res, 46 (2018) 8385–8403.

[31] J. Tian, R. Marino, C. Johnson, J. Locker, Binding of Drug-Activated CAR/Nr1i3 Alters Metabolic Regulation in the Liver, iScience, 9 (2018) 209–228.

[32] H. Tojima, S. Kakizaki, Y. Yamazaki, D. Takizawa, N. Horiguchi, K. Sato, M. Mori, Ligand dependent hepatic gene expression profiles of nuclear receptors CAR and PXR, Toxicol Lett, 212 (2012) 288–297.

[33] F. Al-Salman, N. Plant, Non-coplanar polychlorinated biphenyls (PCBs) are direct agonists for the human pregnane-X receptor and constitutive androstane receptor, and activate target gene expression in a tissue-specific manner, Toxicol Appl Pharmacol, 263 (2012) 7–13.

[34] Y. Chen, Y. Liu, Non-coplanar and coplanar polychlorinated biphenyls potentiate genotoxicity of aflatoxin B1 in a human hepatocyte line by enhancing CYP1A2 and CYP3A4 expression, Environ Pollut, 246 (2019) 945–954.

[35] J. Jin, B. Wahlang, M. Thapa, K.Z. Head, J.E. Hardesty, S. Srivastava, M.L. Merchant, S.N. Rai, R.A. Prough, M.C. Cave, Proteomics and metabolic phenotyping define principal roles for the aryl hydrocarbon receptor in mouse liver, Acta Pharm Sin B, 11 (2021) 3806–3819.

[36] J. Küblbeck, J. Niskanen, P. Honkakoski, Metabolism-Disrupting Chemicals and the Constitutive Androstane Receptor CAR, Cells, 9 (2020).

[37] P. Wei, J. Zhang, M. Egan-Hafley, S. Liang, D.D. Moore, The nuclear receptor CAR mediates specific xenobiotic induction of drug metabolism, Nature, 407 (2000) 920–923.

[38] B. Bhushan, J.W. Stoops, W.M. Mars, A. Orr, W.C. Bowen, S. Paranjpe, G.K. Michalopoulos, TCPOBOP-Induced Hepatomegaly and Hepatocyte Proliferation are Attenuated by Combined Disruption of MET and EGFR Signaling, Hepatology, 69 (2019) 1702–1718.

[39] K. Koral, B. Bhushan, A. Orr, J. Stoops, W.C. Bowen, M.A. Copeland, J. Locker, W.M. Mars, G.K. Michalopoulos, Lymphocyte-Specific Protein-1 Suppresses Xenobiotic-Induced Constitutive Androstane Receptor and Subsequent Yes-Associated Protein-Activated Hepatocyte Proliferation, Am J Pathol, 192 (2022) 887–903.

[40] N.J. Lodato, T. Melia, A. Rampersaud, D.J. Waxman, Sex-Differential Responses of Tumor Promotion-Associated Genes and Dysregulation of Novel Long Noncoding RNAs in Constitutive Androstane Receptor-Activated Mouse Liver, Toxicol Sci, 159 (2017) 25–41.

[41] C.N. Goldfarb, K. Karri, M. Pyatkov, D.J. Waxman, Interplay Between GH-regulated, Sex-biased Liver Transcriptome and Hepatic Zonation Revealed by Single-Nucleus RNA Sequencing, Endocrinology, 163 (2022).

[42] J.L. Dempsey, J.Y. Cui, Regulation of Hepatic Long Noncoding RNAs by Pregnane X Receptor and Constitutive Androstane Receptor Agonists in Mouse Liver, Drug Metab Dispos, 47 (2019) 329–339.

[43] J.Y. Cui, C.D. Klaassen, RNA-Seq reveals common and unique PXR- and CAR-target gene signatures in the mouse liver transcriptome, Biochim Biophys Acta, 1859 (2016) 1198–1217.

[44] W. Huang, J. Zhang, M. Washington, J. Liu, J.M. Parant, G. Lozano, D.D. Moore, Xenobiotic stress induces hepatomegaly and liver tumors via the nuclear receptor constitutive androstane receptor, Mol Endocrinol, 19 (2005) 1646–1653.

[45] B.A. Diwan, R.A. Lubet, J.M. Ward, J.A. Hrabie, J.M. Rice, Tumor-promoting and hepatocarcinogenic effects of 1,4-bis[2-(3,5-dichloropyridyloxy)]benzene (TCPOBOP) in DBA/2NCr and C57BL/6NCr mice and an apparent promoting effect on nasal cavity tumors but not on hepatocellular tumors in F344/NCr rats initiated with N-nitrosodiethylamine, Carcinogenesis, 13 (1992) 1893–1901.

[46] T.A. Dragani, G. Manenti, G. Galliani, G. Della Porta, Promoting effects of 1,4-bis[2-(3,5-dichloropyridyloxy)]benzene in mouse hepatocarcinogenesis, Carcinogenesis, 6 (1985) 225–228.

[47] N. Percie du Sert, A. Ahluwalia, S. Alam, M.T. Avey, M. Baker, W.J. Browne, A. Clark, I.C. Cuthill, U. Dirnagl, M. Emerson, P. Garner, S.T. Holgate, D.W. Howells, V. Hurst, N.A. Karp, S.E. Lazic, K. Lidster, C.J. MacCallum, M. Macleod, E.J. Pearl, O.H. Petersen, F. Rawle, P. Reynolds, K. Rooney, E.S. Sena, S.D. Silberberg, T. Steckler, H. Würbel, Reporting animal research: Explanation and elaboration for the ARRIVE guidelines 2.0, PLoS Biol, 18 (2020) e3000411.

[48] J. Wang, L. Symul, J. Yeung, C. Gobet, J. Sobel, S. Lück, P.O. Westermark, N. Molina, F. Naef, Circadian clock-dependent and -independent posttranscriptional regulation underlies temporal mRNA accumulation in mouse liver, Proc Natl Acad Sci U S A, 115 (2018) E1916–e1925.

[49] C. Droin, J.E. Kholtei, K. Bahar Halpern, C. Hurni, M. Rozenberg, S. Muvkadi, S. Itzkovitz, F. Naef, Space-time logic of liver gene expression at sub-lobular scale, Nat Metab, 3 (2021) 43–58.

[50] T.G. Brooks, A. Manjrekar, A. Mrčela, G.R. Grant, Meta-analysis of Diurnal Transcriptomics in Mouse Liver Reveals Low Repeatability of Rhythm Analyses, J Biol Rhythms, 38 (2023) 556–570.

[51] L.V.M. de Assis, M. Demir, H. Oster, Nonalcoholic Steatohepatitis Disrupts Diurnal Liver Transcriptome Rhythms in Mice, Cell Mol Gastroenterol Hepatol, 16 (2023) 341–354.

[52] A. Poland, I. Mak, E. Glover, R.J. Boatman, F.H. Ebetino, A.S. Kende, 1,4-Bis[2-(3,5-dichloropyridyloxy)]benzene, a potent phenobarbital-like inducer of microsomal monooxygenase activity, Mol Pharmacol, 18 (1980) 571–580.

[53] A. Shin, D.J. Waxman, Impact of Neonatal Activation of Nuclear Receptor CAR (Nr1i3) on Cyp2 Gene Expression in Adult Mouse Liver, Toxicol Sci, 187 (2022) 298–310.

[54] B.T. Sherman, M. Hao, J. Qiu, X. Jiao, M.W. Baseler, H.C. Lane, T. Imamichi, W. Chang, DAVID: a web server for functional enrichment analysis and functional annotation of gene lists (2021 update), Nucleic Acids Res, 50 (2022) W216-221.

[55] X. Wei, M.A. Murphy, N.A. Reddy, Y. Hao, T.G. Eggertsen, J.J. Saucerman, I.M. Bochkis, Redistribution of lamina-associated domains reshapes binding of pioneer factor FOXA2 in development of nonalcoholic fatty liver disease, Genome Res, 32 (2022) 1981–1992.

[56] G. Arguello, E. Balboa, M. Arrese, S. Zanlungo, Recent insights on the role of cholesterol in non-alcoholic fatty liver disease, Biochim Biophys Acta, 1852 (2015) 1765–1778.

[57] R. Gebhardt, A. Baldysiak-Figiel, V. Krügel, E. Ueberham, F. Gaunitz, Hepatocellular expression of glutamine synthetase: an indicator of morphogen actions as master regulators of zonation in adult liver, Prog Histochem Cytochem, 41 (2007) 201–266.

[58] W.H. Lamers, J.L. Vermeulen, T.B. Hakvoort, A.F. Moorman, Expression pattern of glutamine synthetase marks transition from collecting into conducting hepatic veins, J Histochem Cytochem, 47 (1999) 1507–1512.

[59] A. Mardinoglu, M. Uhlen, J. Borén, Broad Views of Non-alcoholic Fatty Liver Disease, Cell Syst, 6 (2018) 7–9.

[60] K. Chella Krishnan, R.R. Floyd, S. Sabir, D.W. Jayasekera, P.V. Leon-Mimila, A.E. Jones, A.A. Cortez, V. Shravah, M. Péterfy, L. Stiles, S. Canizales-Quinteros, A.S. Divakaruni, A. Huertas-Vazquez, A.J. Lusis, Liver Pyruvate Kinase Promotes NAFLD/NASH in Both Mice and Humans in a Sex-Specific Manner, Cell Mol Gastroenterol Hepatol, 11 (2021) 389–406.

[61] A. Cherubini, M. Ostadreza, O. Jamialahmadi, S. Pelusi, E. Rrapaj, E. Casirati, G. Passignani, M. Norouziesfahani, E. Sinopoli, G. Baselli, C. Meda, P. Dongiovanni, D. Dondossola, N. Youngson, A. Tourna, S. Chokshi, E. Bugianesi, S. Della Torre, D. Prati, S. Romeo, L. Valenti, Interaction between estrogen receptor-α and PNPLA3 p.I148M variant drives fatty liver disease susceptibility in women, Nat Med, 29 (2023) 2643–2655.

[62] E. Ericson, L. Bergenholm, A.C. Andréasson, C.I. Dix, J. Knöchel, S.F. Hansson, R. Lee, J. Schumi, M. Antonsson, O. Fjellström, P. Nasr, M. Liljeblad, B. Carlsson, S. Kechagias, D. Lindén, M. Ekstedt, Hepatic patatin-like phospholipase domain-containing 3 levels are increased in I148M risk allele carriers and correlate with NAFLD in humans, Hepatol Commun, 6 (2022) 2689–2701.

[63] A. Cherubini, E. Casirati, M. Tomasi, L. Valenti, PNPLA3 as a therapeutic target for fatty liver disease: the evidence to date, Expert Opin Ther Targets, 25 (2021) 1033–1043.

[64] K. Kumagai, K. Tabu, F. Sasaki, Y. Takami, Y. Morinaga, S. Mawatari, S. Hashimoto, S. Tanoue, S. Kanmura, T. Tamai, A. Moriuchi, H. Uto, H. Tsubouchi, A. Ido, Glycoprotein Nonmetastatic Melanoma B (Gpnmb)-Positive Macrophages Contribute to the Balance between Fibrosis and Fibrolysis during the Repair of Acute Liver Injury in Mice, PLoS One, 10 (2015) e0143413.

[65] A. Naim, Q. Pan, M.S. Baig, Matrix Metalloproteinases (MMPs) in Liver Diseases, J Clin Exp Hepatol, 7 (2017) 367–372.

[66] H.P. Ma, H.L. Chang, O.A. Bamodu, V.K. Yadav, T.Y. Huang, A.T.H. Wu, C.T. Yeh, S.H. Tsai, W.H. Lee, Collagen 1A1 (COL1A1) Is a Reliable Biomarker and Putative Therapeutic Target for Hepatocellular Carcinogenesis and Metastasis, Cancers (Basel), 11 (2019).

[67] S. Cazanave, A. Podtelezhnikov, K. Jensen, M. Seneshaw, D.P. Kumar, H.K. Min, P.K. Santhekadur, B. Banini, A.G. Mauro, M.O. A R. Vincent, K.Q. Tanis, A.L. Webber, L. Wang, P. Bedossa, F. Mirshahi, A.J. Sanyal, The Transcriptomic Signature Of Disease Development And Progression Of Nonalcoholic Fatty Liver Disease, Sci Rep, 7 (2017) 17193.

[68] T. Kawamoto, S. Kakizaki, K. Yoshinari, M. Negishi, Estrogen activation of the nuclear orphan receptor CAR (constitutive active receptor) in induction of the mouse Cyp2b10 gene, Mol Endocrinol, 14 (2000) 1897–1905.

[69] G.M. Ledda-Columbano, M. Pibiri, D. Concas, F. Molotzu, G. Simbula, C. Cossu, A. Columbano, Sex difference in the proliferative response of mouse hepatocytes to treatment with the CAR ligand, TCPOBOP, Carcinogenesis, 24 (2003) 1059–1065.

[70] J. Ren, Z. Chen, W. Zhang, L. Li, R. Sun, C. Deng, Z. Fei, Z. Sheng, L. Wang, X. Sun, Z. Wang, J. Fei, Increased fat mass and insulin resistance in mice lacking pancreatic lipase-related protein 1, J Nutr Biochem, 22 (2011) 691–698.

[71] F. Wagner, I. Ruf, T. Lehmann, R. Hofmann, S. Ortmann, C. Schiffmann, M. Hiller, C. Stefen, H. Stuckas, Reconstruction of evolutionary changes in fat and toxin consumption reveals associations with gene losses in mammals: A case study for the lipase inhibitor PNLIPRP1 and the xenobiotic receptor NR1I3, J Evol Biol, 35 (2022) 225–239.

[72] J. Wang, H. Yu, W. Dong, C. Zhang, M. Hu, W. Ma, X. Jiang, H. Li, P. Yang, D. Xiang, N6-Methyladenosine-Mediated Up-Regulation of FZD10 Regulates Liver Cancer Stem Cells’ Properties and Lenvatinib Resistance Through WNT/β-Catenin and Hippo Signaling Pathways, Gastroenterology, 164 (2023) 990–1005.

[73] M. Arrese, D. Cabrera, A.M. Kalergis, A.E. Feldstein, Innate Immunity and Inflammation in NAFLD/NASH, Dig Dis Sci, 61 (2016) 1294–1303.

[74] H. Tilg, T.E. Adolph, M. Dudek, P. Knolle, Non-alcoholic fatty liver disease: the interplay between metabolism, microbes and immunity, Nat Metab, 3 (2021) 1596–1607.

[75] M. Vacca, J. Leslie, S. Virtue, B.Y.H. Lam, O. Govaere, D. Tiniakos, S. Snow, S. Davies, K. Petkevicius, Z. Tong, V. Peirce, M.J. Nielsen, Z. Ament, W. Li, T. Kostrzewski, D.J. Leeming, V. Ratziu, M.E.D. Allison, Q.M. Anstee, J.L. Griffin, F. Oakley, A. Vidal-Puig, Bone morphogenetic protein 8B promotes the progression of non-alcoholic steatohepatitis, Nat Metab, 2 (2020) 514–531.

[76] N. Dali-Youcef, M. Vix, F. Costantino, H. El-Saghire, B. Lhermitte, C. Callari, J. D’Agostino, S. Perretta, S. Paveliu, M. Gualtierotti, E. Dumeny, M.A. Oudot, A. Jaulin, D. Dembélé, M.B. Zeisel, C. Tomasetto, T.F. Baumert, M. Doffoël, Interleukin-32 Contributes to Human Nonalcoholic Fatty Liver Disease and Insulin Resistance, Hepatol Commun, 3 (2019) 1205–1220.

[77] J. Oliva, F. Bardag-Gorce, B.A. French, J. Li, S.W. French, Independent phenotype of binuclear hepatocytes and cellular localization of UbD, Exp Mol Pathol, 89 (2010) 103–108.

[78] L.A. Navarro, A. Wree, D. Povero, M.P. Berk, A. Eguchi, S. Ghosh, B.G. Papouchado, S.C. Erzurum, A.E. Feldstein, Arginase 2 deficiency results in spontaneous steatohepatitis: a novel link between innate immune activation and hepatic de novo lipogenesis, J Hepatol, 62 (2015) 412–420.

[79] J. Gu, Y. Zhang, D. Xu, Z. Zhao, Y. Zhang, Y. Pan, P. Cao, Z. Wang, Y. Chen, Ethanol-induced hepatic steatosis is modulated by glycogen level in the liver, J Lipid Res, 56 (2015) 1329–1339.

[80] X. Xiong, H. Kuang, S. Ansari, T. Liu, J. Gong, S. Wang, X.Y. Zhao, Y. Ji, C. Li, L. Guo, L. Zhou, Z. Chen, P. Leon-Mimila, M.T. Chung, K. Kurabayashi, J. Opp, F. Campos-Pérez, H. Villamil-Ramírez, S. Canizales-Quinteros, R. Lyons, C.N. Lumeng, B. Zhou, L. Qi, A. Huertas-Vazquez, A.J. Lusis, X.Z.S. Xu, S. Li, Y. Yu, J.Z. Li, J.D. Lin, Landscape of Intercellular Crosstalk in Healthy and NASH Liver Revealed by Single-Cell Secretome Gene Analysis, Mol Cell, 75 (2019) 644–660.e645.

[81] Y. Tsuchiya, T. Seki, K. Kobayashi, S. Komazawa-Sakon, S. Shichino, T. Nishina, K. Fukuhara, K. Ikejima, H. Nagai, Y. Igarashi, S. Ueha, A. Oikawa, S. Tsurusaki, S. Yamazaki, C. Nishiyama, T. Mikami, H. Yagita, K. Okumura, T. Kido, A. Miyajima, K. Matsushima, M. Imasaka, K. Araki, T. Imamura, M. Ohmuraya, M. Tanaka, H. Nakano, Fibroblast growth factor 18 stimulates the proliferation of hepatic stellate cells, thereby inducing liver fibrosis, Nat Commun, 14 (2023) 6304.

[82] J.Y. Cha, J.J. Repa, The liver X receptor (LXR) and hepatic lipogenesis. The carbohydrate-response element-binding protein is a target gene of LXR, J Biol Chem, 282 (2007) 743–751.

[83] J.P. Rooney, K. Oshida, R. Kumar, W.S. Baldwin, J.C. Corton, Chemical Activation of the Constitutive Androstane Receptor Leads to Activation of Oxidant-Induced Nrf2, Toxicol Sci, 167 (2019) 172–189.

[84] A.A. Yarushkin, M.E. Mazin, Y.A. Pustylnyak, E.A. Prokopyeva, V.O. Pustylnyak, Activation of the Akt pathway by a constitutive androstane receptor agonist results in β-catenin activation, Eur J Pharmacol, 879 (2020) 173135.

[85] K.H. Kim, J.M. Choi, F. Li, A. Arizpe, C.R. Wooton-Kee, S. Anakk, S.Y. Jung, M.J. Finegold, D.D. Moore, Xenobiotic Nuclear Receptor Signaling Determines Molecular Pathogenesis of Progressive Familial Intrahepatic Cholestasis, Endocrinology, 159 (2018) 2435–2446.

[86] P. Hao, D.J. Waxman, STAT5 Regulation of Sex-Dependent Hepatic CpG Methylation at Distal Regulatory Elements Mapping to Sex-Biased Genes, Mol Cell Biol, 41 (2021).

[87] M.G. Holloway, Y. Cui, E.V. Laz, A. Hosui, L. Hennighausen, D.J. Waxman, Loss of sexually dimorphic liver gene expression upon hepatocyte-specific deletion of Stat5a-Stat5b locus, Endocrinology, 148 (2007) 1977–1986.

[88] M. Grohmann, F. Wiede, G.T. Dodd, E.N. Gurzov, G.J. Ooi, T. Butt, A.A. Rasmiena, S. Kaur, T. Gulati, P.K. Goh, A.E. Treloar, S. Archer, W.A. Brown, M. Muller, M.J. Watt, O. Ohara, C.A. McLean, T. Tiganis, Obesity Drives STAT-1-Dependent NASH and STAT-3-Dependent HCC, Cell, 175 (2018) 1289–1306.e1220.

[89] F. Wang, X. Zhang, W. Liu, Y. Zhou, W. Wei, D. Liu, C.C. Wong, J.J.Y. Sung, J. Yu, Activated Natural Killer Cell Promotes Nonalcoholic Steatohepatitis Through Mediating JAK/STAT Pathway, Cell Mol Gastroenterol Hepatol, 13 (2022) 257–274.

[90] A. Mohs, N. Kuttkat, T. Otto, S.A. Youssef, A. De Bruin, C. Trautwein, MyD88-dependent signaling in non-parenchymal cells promotes liver carcinogenesis, Carcinogenesis, 41 (2020) 171–181.

[91] Y. Liu, H. Chen, X. Yan, J. Zhang, Z. Deng, M. Huang, J. Gu, J. Zhang, MyD88 in myofibroblasts enhances nonalcoholic fatty liver disease-related hepatocarcinogenesis via promoting macrophage M2 polarization, Cell Commun Signal, 22 (2024) 86.

[92] K. Neumann, B. Schiller, G. Tiegs, NLRP3 Inflammasome and IL-33: Novel Players in Sterile Liver Inflammation, Int J Mol Sci, 19 (2018).

[93] Z. Chen, R. Yu, Y. Xiong, F. Du, S. Zhu, A vicious circle between insulin resistance and inflammation in nonalcoholic fatty liver disease, Lipids Health Dis, 16 (2017) 203.

[94] T.T. Zhang, Y. Wang, X.W. Zhang, K.Y. Yang, X.Q. Miao, G.H. Zhao, MiR-200c-3p Regulates DUSP1/MAPK Pathway in the Nonalcoholic Fatty Liver After Laparoscopic Sleeve Gastrectomy, Front Endocrinol (Lausanne), 13 (2022) 792439.

[95] S. Aggarwal, N. Trehanpati, P. Nagarajan, G. Ramakrishna, The Clock-NAD(+) -Sirtuin connection in nonalcoholic fatty liver disease, J Cell Physiol, (2022).

[96] E. de Gregorio, A. Colell, A. Morales, M. Marí, Relevance of SIRT1-NF-κB Axis as Therapeutic Target to Ameliorate Inflammation in Liver Disease, Int J Mol Sci, 21 (2020).

[97] M. Sakai, Exploring the signal-dependent transcriptional regulation involved in the liver pathology of type 2 diabetes, Diabetol Int, 14 (2023) 15–20.

[98] R. Chen, J. Du, H. Zhu, Q. Ling, The role of cGAS-STING signalling in liver diseases, JHEP Rep, 3 (2021) 100324.

[99] R. Donne, M. Saroul-Ainama, P. Cordier, A. Hammoutene, C. Kabore, M. Stadler, I. Nemazanyy, I. Galy-Fauroux, M. Herrag, T. Riedl, M. Chansel-Da Cruz, S. Caruso, S. Bonnafous, R. Öllinger, R. Rad, K. Unger, A. Tran, J.P. Couty, P. Gual, V. Paradis, S. Celton-Morizur, M. Heikenwalder, P. Revy, C. Desdouets, Replication stress triggered by nucleotide pool imbalance drives DNA damage and cGAS-STING pathway activation in NAFLD, Dev Cell, 57 (2022) 1728–1741.e1726.

[100] Y. Liu, F. Yu, Y. Han, Q. Li, Z. Cao, X. Xiang, S. Jiang, X. Wang, J. Lu, R. Lai, H. Wang, W. Cai, S. Bao, Q. Xie, SUMO-specific protease 3 is a key regulator for hepatic lipid metabolism in non-alcoholic fatty liver disease, Sci Rep, 6 (2016) 37351.

[101] S. Zang, X. Ma, Y. Wu, W. Liu, H. Cheng, J. Li, J. Liu, A. Huang, PGE(2) synthesis and signaling in malignant transformation and progression of human hepatocellular carcinoma, Hum Pathol, 63 (2017) 120–127.

[102] Y. Bai, F. Xie, F. Miao, J. Long, S. Huang, H. Huang, J. Lin, D. Wang, X. Yang, J. Bian, J. Mao, X. Wang, Y. Mao, X. Sang, H. Zhao, The diagnostic and prognostic role of RhoA in hepatocellular carcinoma, Aging (Albany NY), 11 (2019) 5158–5172.

[103] J. Zan, X. Zhao, X. Deng, H. Ding, B. Wang, M. Lu, Z. Wei, Z. Huang, S. Wang, Paraspeckle Promotes Hepatocellular Carcinoma Immune Escape by Sequestering IFNGR1 mRNA, Cell Mol Gastroenterol Hepatol, 12 (2021) 465–487.

[104] M. Shen, R. Zhang, W. Jia, Z. Zhu, X. Zhao, L. Zhao, G. Huang, J. Liu, Nuclear scaffold protein p54(nrb)/NONO facilitates the hypoxia-enhanced progression of hepatocellular carcinoma, Oncogene, 40 (2021) 4167–4183.

[105] H. Wu, J. Zhang, Y. Bai, S. Zhang, Z. Zhang, W. Tong, P. Han, B. Fu, Y. Zhang, Z. Shen, DCP1A is an unfavorable prognostic-related enhancer RNA in hepatocellular carcinoma, Aging (Albany NY), 13 (2021) 23020–23035.

[106] B. Shu, Y.X. Zhou, H. Li, R.Z. Zhang, C. He, X. Yang, The METTL3/MALAT1/PTBP1/USP8/TAK1 axis promotes pyroptosis and M1 polarization of macrophages and contributes to liver fibrosis, Cell Death Discov, 7 (2021) 368.

[107] Y. Zhu, J. Xu, W. Hu, F. Wang, Y. Zhou, W. Gong, W. Xu, Inhibiting USP8 overcomes hepatocellular carcinoma resistance via suppressing receptor tyrosine kinases, Aging (Albany NY), 13 (2021) 14999–15012.

[108] M.J. Kim, B. Choi, J.Y. Kim, Y. Min, D.H. Kwon, J. Son, J.S. Lee, J.S. Lee, E. Chun, K.Y. Lee, USP8 regulates liver cancer progression via the inhibition of TRAF6-mediated signal for NF-κB activation and autophagy induction by TLR4, Transl Oncol, 15 (2022) 101250.

[109] A. Takahashi, T.M. Loo, R. Okada, F. Kamachi, Y. Watanabe, M. Wakita, S. Watanabe, S. Kawamoto, K. Miyata, G.N. Barber, N. Ohtani, E. Hara, Downregulation of cytoplasmic DNases is implicated in cytoplasmic DNA accumulation and SASP in senescent cells, Nat Commun, 9 (2018) 1249.

[110] M.K. Atianand, W. Hu, A.T. Satpathy, Y. Shen, E.P. Ricci, J.R. Alvarez-Dominguez, A. Bhatta, S.A. Schattgen, J.D. McGowan, J. Blin, J.E. Braun, P. Gandhi, M.J. Moore, H.Y. Chang, H.F. Lodish, D.R. Caffrey, K.A. Fitzgerald, A Long Noncoding RNA lincRNA-EPS Acts as a Transcriptional Brake to Restrain Inflammation, Cell, 165 (2016) 1672–1685.

[111] C.Y. Yang, H.C. Chuang, C.Y. Tsai, Y.Z. Xiao, J.Y. Yang, R.H. Huang, Y.C. Shih, T.H. Tan, DUSP11 Attenuates Lipopolysaccharide-Induced Macrophage Activation by Targeting TAK1, J Immunol, 205 (2020) 1644–1652.

[112] R. Raghu, B. Jesudas, G. Bhavani, D. Ezhilarasan, S. Karthikeyan, Silibinin mitigates zidovudine-induced hepatocellular degenerative changes, oxidative stress and hyperlipidaemia in rats, Hum Exp Toxicol, 34 (2015) 1031–1042.

[113] G. Baffy, Origins of Portal Hypertension in Nonalcoholic Fatty Liver Disease, Dig Dis Sci, 63 (2018) 563–576.

[114] K. Nishi, H. Yagi, M. Ohtomo, S. Nagata, D. Udagawa, T. Tsuchida, T. Morisaku, Y. Kitagawa, A thioacetamide-induced liver fibrosis model for pre-clinical studies in microminipig, Sci Rep, 13 (2023) 14996.

[115] R. Huang, J. Deng, C.P. Zhu, S.Q. Liu, Y.L. Cui, F. Chen, X. Zhang, X. Tao, W.F. Xie, Sulodexide attenuates liver fibrosis in mice by restoration of differentiated liver sinusoidal endothelial cell, Biomed Pharmacother, 160 (2023) 114396.

[116] H. Malhi, R.J. Kaufman, Endoplasmic reticulum stress in liver disease, J Hepatol, 54 (2011) 795–809.

[117] J. Li, X. Li, D. Liu, S. Zhang, N. Tan, H. Yokota, P. Zhang, Phosphorylation of eIF2α signaling pathway attenuates obesity-induced non-alcoholic fatty liver disease in an ER stress and autophagy-dependent manner, Cell Death Dis, 11 (2020) 1069.

[118] S.A. Dabravolski, E.E. Bezsonov, A.N. Orekhov, The role of mitochondria dysfunction and hepatic senescence in NAFLD development and progression, Biomed Pharmacother, 142 (2021) 112041.

[119] B. Fromenty, M. Roden, Mitochondrial alterations in fatty liver diseases, J Hepatol, 78 (2023) 415–429.

[120] H. Wang, Y. Liu, D. Wang, Y. Xu, R. Dong, Y. Yang, Q. Lv, X. Chen, Z. Zhang, The Upstream Pathway of mTOR-Mediated Autophagy in Liver Diseases, Cells, 8 (2019).

[121] M. Régnier, T. Carbinatti, L. Parlati, F. Benhamed, C. Postic, The role of ChREBP in carbohydrate sensing and NAFLD development, Nat Rev Endocrinol, 19 (2023) 336–349.

[122] T. Carbinatti, M. Régnier, L. Parlati, F. Benhamed, C. Postic, New insights into the inter-organ crosstalk mediated by ChREBP, Front Endocrinol (Lausanne), 14 (2023) 1095440.

[123] W. Li, C.C. Wong, X. Zhang, W. Kang, G. Nakatsu, Q. Zhao, H. Chen, M.Y.Y. Go, P.W.Y. Chiu, X. Wang, J. Ji, X. Li, Z. Cai, E.K.W. Ng, J. Yu, CAB39L elicited an anti-Warburg effect via a LKB1-AMPK-PGC1α axis to inhibit gastric tumorigenesis, Oncogene, 37 (2018) 6383–6398.

[124] T. Luo, S. Yang, T. Zhao, H. Zhu, C. Chen, X. Shi, D. Chen, K. Wang, K. Jiang, D. Xu, M. Cheng, J. Li, W. Li, W. Xu, L. Zhou, M. Jiang, B. Xu, Hepatocyte DDX3X protects against drug-induced acute liver injury via controlling stress granule formation and oxidative stress, Cell Death Dis, 14 (2023) 400.

[125] J. Feng, S. Qiu, S. Zhou, Y. Tan, Y. Bai, H. Cao, J. Guo, Z. Su, mTOR: A Potential New Target in Nonalcoholic Fatty Liver Disease, Int J Mol Sci, 23 (2022).

[126] M. Yang, Y. Lu, W. Piao, H. Jin, The Translational Regulation in mTOR Pathway, Biomolecules, 12 (2022).

[127] S. Fang, L. Zheng, L. Shen, Y. Su, J. Ding, W. Chen, X. Chen, W. Chen, G. Shu, M. Chen, Z. Zhao, J. Tu, J. Ji, Inactivation of KDM5A suppresses growth and enhances chemosensitivity in liver cancer by modulating ROCK1/PTEN/AKT pathway, Eur J Pharmacol, 940 (2023) 175465.

[128] R.R. Maronpot, K. Yoshizawa, A. Nyska, T. Harada, G. Flake, G. Mueller, B. Singh, J.M. Ward, Hepatic enzyme induction: histopathology, Toxicol Pathol, 38 (2010) 776–795.

[129] L. Deng, S. Kersten, R. Stienstra, Triacylglycerol uptake and handling by macrophages: From fatty acids to lipoproteins, Prog Lipid Res, 92 (2023) 101250.

[130] S. Lefere, F. Tacke, Macrophages in obesity and non-alcoholic fatty liver disease: Crosstalk with metabolism, JHEP Rep, 1 (2019) 30–43.

[131] S. Ministrini, F. Montecucco, A. Sahebkar, F. Carbone, Macrophages in the pathophysiology of NAFLD: The role of sex differences, Eur J Clin Invest, 50 (2020) e13236.

[132] A. Marmugi, C. Lukowicz, F. Lasserre, A. Montagner, A. Polizzi, S. Ducheix, A. Goron, L. Gamet-Payrastre, S. Gerbal-Chaloin, J.M. Pascussi, M. Moldes, T. Pineau, H. Guillou, L. Mselli-Lakhal, Activation of the Constitutive Androstane Receptor induces hepatic lipogenesis and regulates Pnpla3 gene expression in a LXR-independent way, Toxicol Appl Pharmacol, 303 (2016) 90–100.

[133] L. Coassolo, T. Liu, Y. Jung, N.P. Taylor, M. Zhao, G.W. Charville, S.B. Nissen, H. Yki-Jarvinen, R.B. Altman, K.J. Svensson, Mapping transcriptional heterogeneity and metabolic networks in fatty livers at single-cell resolution, iScience, 26 (2023) 105802.

[134] B. Dong, P.K. Saha, W. Huang, W. Chen, L.A. Abu-Elheiga, S.J. Wakil, R.D. Stevens, O. Ilkayeva, C.B. Newgard, L. Chan, D.D. Moore, Activation of nuclear receptor CAR ameliorates diabetes and fatty liver disease, Proceedings of the National Academy of Sciences of the United States of America, 106 (2009) 18831–18836.

[135] J. Gao, J. He, Y. Zhai, T. Wada, W. Xie, The constitutive androstane receptor is an anti-obesity nuclear receptor that improves insulin sensitivity, J Biol Chem, 284 (2009) 25984–25992.

[136] J. Tian, J. Locker, Gadd45 in the Liver: Signal Transduction and Transcriptional Mechanisms, Adv Exp Med Biol, 1360 (2022) 87–99.

[137] J. Miao, S. Fang, Y. Bae, J.K. Kemper, Functional inhibitory cross-talk between constitutive androstane receptor and hepatic nuclear factor-4 in hepatic lipid/glucose metabolism is mediated by competition for binding to the DR1 motif and to the common coactivators, GRIP-1 and PGC-1alpha, J Biol Chem, 281 (2006) 14537–14546.

[138] E.M. Kachaylo, A.A. Yarushkin, V.O. Pustylnyak, Constitutive androstane receptor activation by 2,4,6-triphenyldioxane-1,3 suppresses the expression of the gluconeogenic genes, Eur J Pharmacol, 679 (2012) 139–143.

[139] J.M. Maglich, D.C. Lobe, J.T. Moore, The nuclear receptor CAR (NR1I3) regulates serum triglyceride levels under conditions of metabolic stress, J Lipid Res, 50 (2009) 439–445.

[140] J. Gao, J. Yan, M. Xu, S. Ren, W. Xie, CAR Suppresses Hepatic Gluconeogenesis by Facilitating the Ubiquitination and Degradation of PGC1α, Mol Endocrinol, 29 (2015) 1558–1570.

[141] M. Martin-Grau, D. Monleon, Sex dimorphism and metabolic profiles in management of metabolic-associated fatty liver disease, World J Clin Cases, 11 (2023) 1236–1244.

[142] S. Smati, A. Polizzi, A. Fougerat, S. Ellero-Simatos, Y. Blum, Y. Lippi, M. Régnier, A. Laroyenne, M. Huillet, M. Arif, C. Zhang, F. Lasserre, A. Marrot, T. Al Saati, J. Wan, C. Sommer, C. Naylies, A. Batut, C. Lukowicz, T. Fougeray, B. Tramunt, P. Dubot, L. Smith, J. Bertrand-Michel, N. Hennuyer, J.P. Pradere, B. Staels, R. Burcelin, F. Lenfant, J.F. Arnal, T. Levade, L. Gamet-Payrastre, S. Lagarrigue, N. Loiseau, S. Lotersztajn, C. Postic, W. Wahli, C. Bureau, M. Guillaume, A. Mardinoglu, A. Montagner, P. Gourdy, H. Guillou, Integrative study of diet-induced mouse models of NAFLD identifies PPARα as a sexually dimorphic drug target, Gut, 71 (2022) 807–821.

[143] D.J. Waxman, M.G. Holloway, Sex differences in the expression of hepatic drug metabolizing enzymes, Mol Pharmacol, 76 (2009) 215–228.

[144] J.B. Steinman, M.A. Salomao, U.B. Pajvani, Zonation in NASH - A key paradigm for understanding pathophysiology and clinical outcomes, Liver Int, 41 (2021) 2534–2546.

